# Genetic architecture of the red blood cell proteome in genetically diverse mice reveals central role of hemoglobin beta cysteine redox status in maintaining circulating glutathione pools

**DOI:** 10.1101/2025.02.27.640676

**Authors:** Gregory R. Keele, Monika Dzieciatkowska, Ariel M. Hay, Matthew Vincent, Callan O’Connor, Daniel Stephenson, Julie A. Reisz, Travis Nemkov, Kirk C. Hansen, Grier P. Page, James C Zimring, Gary A. Churchill, Angelo D’Alessandro

**Author notes:** **Corresponding authors:** Gary A. Churchill, PhD The Jackson Laboratory Bar Harbor, ME, USA., James C Zimring, MD PhD Department of Pathology University of Virginia 200 Jeanette Lancaster Way Charlottesville, VA 22903, Angelo D’Alessandro, PhD ***lead contact** Department of Biochemistry and Molecular Genetics University of Colorado Anschutz Medical Campus 12801 East 17th Ave., Aurora, CO 80045 www.dalessandrolab.com.

## Abstract

Red blood cells (RBCs) transport oxygen but accumulate oxidative damage over time, reducing function in vivo and during storage—critical for transfusions. To explore genetic influences on RBC resilience, we profiled proteins, metabolites, and lipids from fresh and stored RBCs obtained from 350 genetically diverse mice. Our analysis identified over 6,000 quantitative trait loci (QTL). Compared to other tissues, prevalence of *trans* genetic effects over *cis* reflects the absence of *de novo* protein synthesis in anucleated RBCs. QTL hotspots at Hbb, Hba, Mon1a, and storage-specific Steap3 linked ferroptosis to hemolysis. Proteasome components clustered at multiple loci, underscoring the importance of degrading oxidized proteins. Post-translational modifications (PTMs) mapped predominantly to hemoglobins, particularly cysteine residues. Loss of reactive C93 in humanized mice (HBB C93A) disrupted redox balance, affecting glutathione pools, protein glutathionylation, and redox PTMs. These findings highlight genetic regulation of RBC oxidation, with implications for transfusion biology and oxidative stress-dependent hemolytic disorders.

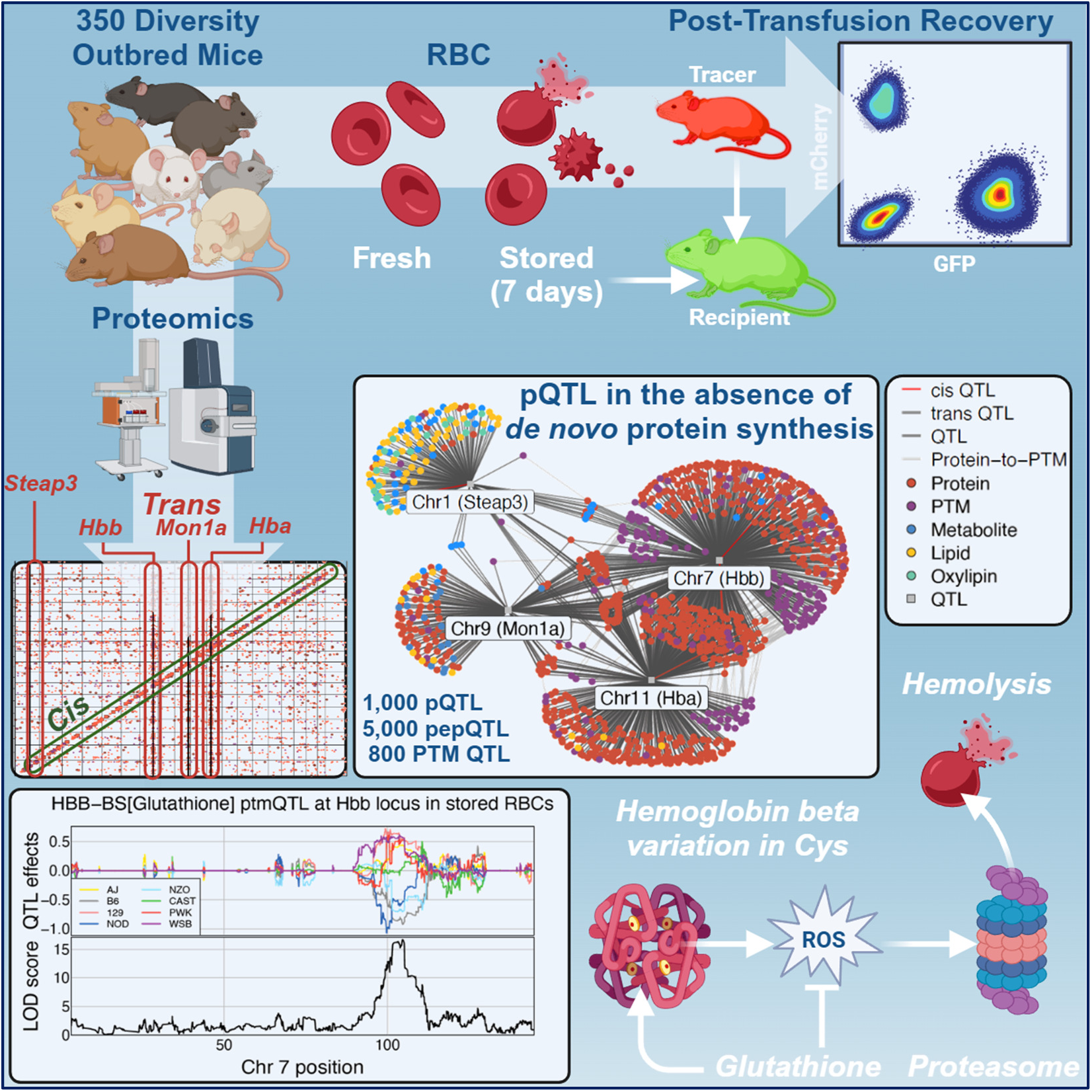

## INTRODUCTION

Red blood cells (RBCs) constitute 99% of the cells in the circulatory system and approximately 83% of the total cells in an adult human^1^. Due to their role in oxygen transport, RBCs have evolved a relatively simple proteome. During maturation, erythroid precursors lose nuclei and organelles, along with their capacity to synthesize new proteins to replace damaged components^2^. Mature RBCs contain about 250- 270 million multimeric hemoglobin molecules per cell, accounting for roughly 98% of the total cytosolic proteins^3^. Nevertheless, proteomics studies over the past two decades have identified around 4,600 distinct RBC proteins^4^, including at least 77 active transporters^5^ that scavenge and release small molecules into the bloodstream, fulfilling critical secondary functions as RBCs circulate throughout the body. As such, the RBC proteome provides insight into proteome-metabolome interactions essential to understanding systems physiology and organ function, a founding principle in clinical biochemistry.

RBCs are exposed to extreme oxidative stress as they age in circulation, traveling from lung alveoli with 100 mmHg pO2 to peripheral capillaries with as low as 20-40 mmHg pO_2_. To facilitate oxygen transport, RBCs carry about 66% of bodily iron in the prosthetic group of hemoglobin^6^, an evolutionary adaptation that promotes iron-dependent Fenton and Haber-Weiss chemistry, generating 1-2 µmol/s of superoxide per liter of packed RBCs due to hemoglobin auto-oxidation, occurring at an estimated rate of 0.5-3% of hemoglobin per day^7^. These rates are exacerbated with aging, high-altitude, pathological hypoxia, exercise, or medical interventions such as chemotherapy^8^.

Understanding RBC responses to oxidative stress is crucial for improving the quality of stored blood products in transfusion medicine^9^. RBC transfusion is the most common hospital medical procedure after vaccination, a life-saving intervention for over five million Americans annually, and for an estimated one billion people since its invention over a century ago^10^. Refrigerated storage is necessary to preserve blood until needed for transfusion therapies, but it leads to the progressive accumulation of biochemical and morphological alterations, mainly triggered by oxidative stress to proteins and lipids, a phenomenon known as the storage lesion^9^. Murine models of blood storage have identified genetic factors that contribute to the altered capacity of stored RBCs (*e.g.,* the ferrireductase STEAP3)^11,12^ and mitigate oxidative damage (*e.g.,* activation of the pentose phosphate pathway by G6PD to generate reducing equivalents)^13–15^. These factors are key drivers of the storage lesion and post-transfusion recovery (PTR), the capacity of end-of-storage RBCs to circulate 24 hours after transfusion. In humans, metabolic lesions (especially lipid peroxidation) and the genetic alterations underpinning them have been linked to poor hemoglobin increments in transfusion recipients^12,16–18^, a measurement of transfusion efficacy that indicates increases in clinical measurements of circulating hemoglobin levels in patients requiring transfusions.

In this study, we aimed to elucidate the genetic basis of variation in the RBC proteome and its response to refrigerated storage. We obtained fresh and stored murine RBCs from genetically diverse mice and used mass spectrometry to characterize the proteome. We identified proteins and their post-translational modifications (PTMs) that change in response to storage. Genetic mapping revealed over 1,000 protein QTL (pQTL), approximately 5,000 peptide QTL (pepQTL), and around 800 PTM QTL (ptmQTL). Our results reveal for the first time that the regulation of the murine RBC proteome is dominated by *trans*-regulatory genetic variation at the hemoglobin alpha and beta loci on chromosomes 11 and 7, respectively. Specifically, the *trans* effects at hemoglobin beta stem from a variant that introduces an extra cysteine in the HBB sequence (Cys13). This variant, along with a polymorphic variants in the ferrireductase STEAP3 on chromosome 1 impact redox (glutathione and oxidative PTM) homeostasis in fresh and stored RBCs, delinneating protein markers of the storage lesion and protein, peptide, and PTM correlates to PTR, a marker of transfusion efficacy. Using a humanized mouse model of *Hbb*, we confirmed that perturbing the total numbers of HBB cysteine residues (Cys93) resulted in altered glutathione pools, redox homeostasis, and the compensatory up-regulation of proteolytic machinery to remove redox damaged components.

## RESULTS

### Profiling the murine red blood cell (RBC) proteome

We analyzed the proteomes of RBCs from 350 Diversity Outbred (J:DO) mice obtained from the Jackson Laboratory^19^ using data-independent acquisition (DIA) mass spectrometry (**Figure 1A**). The J:DO mice are descended from eight inbred founder strains: A/J (AJ), C57BL/6J (B6), 129S1/SvlmJ (129), NOD/ShiLtJ (NOD), NZO/HILtJ (NZO), CAST/EiJ (CAST), PWK/PhJ (PWK), and WSB/EiJ (WSB), that capture a broad sampling of genetic diversity across the *Mus musculus* species^20,21^. For each J:DO mouse, we obtained samples of fresh and stored RBCs for analysis. Stored RBCs were refrigerated for seven days, approximating end-of-storage human packed RBCs^11^. We quantified the abundance of 18,285 peptides, representing 2,134 proteins, and 7,537 peptides with any of ten different PTMs, including oxidation (M, C), deamidation followed by methylation (N, Q), methylation (K, D, E), phosphorylation (S, T, Y), acetylation (K), irreversible beta elimination of C thiol to dehydroalanine (Cys→Dha), glutathionylation (C), lactylation (K), and dioxidation of C (sulfonic acid or sulfenic acid).

**Figure 1.**
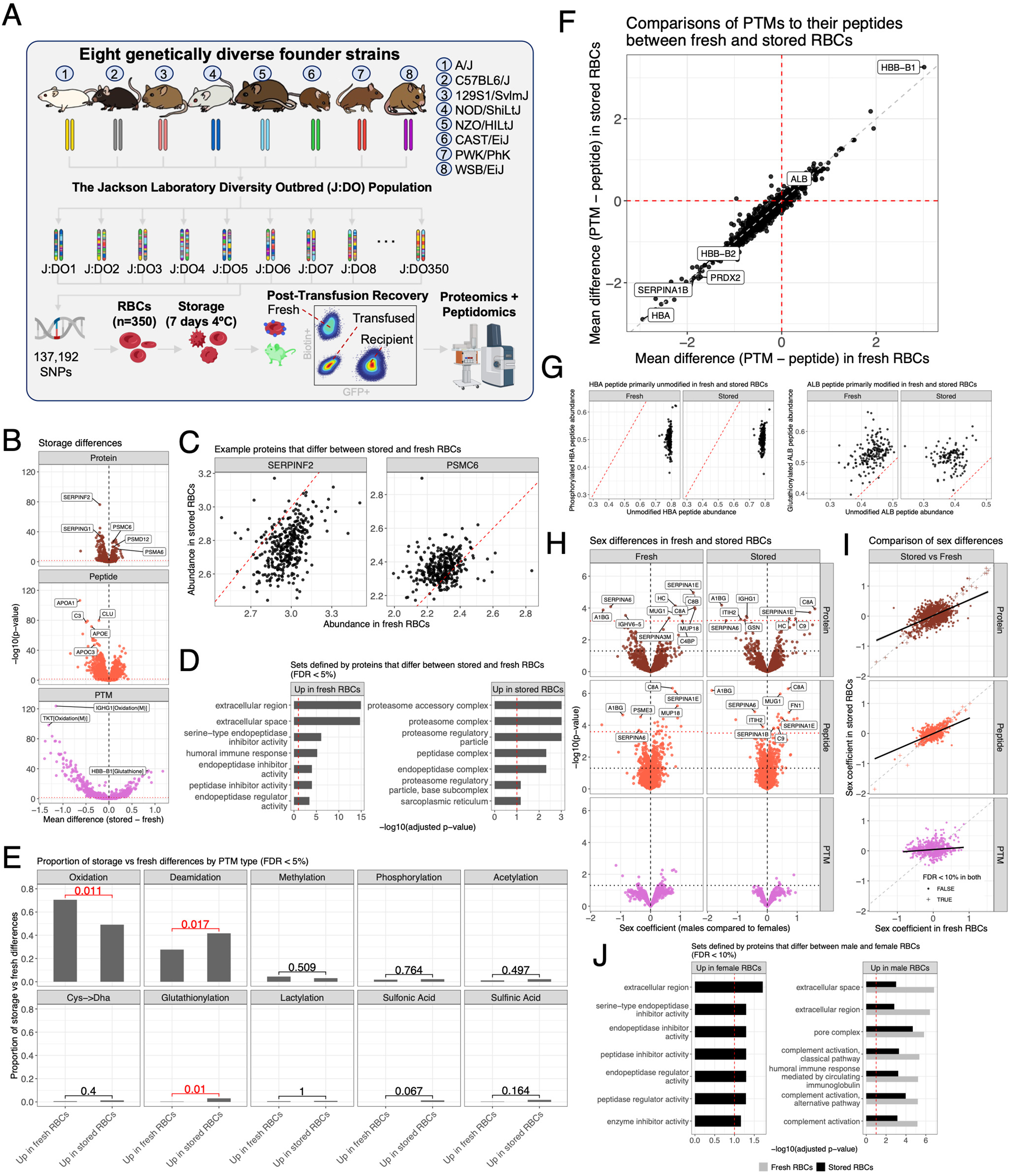
Storage and sex dynamics in a genetically diverse murine RBC proteome. (A) Diagram of J:DO resource population and the experimental approach to profiling the proteomes of fresh and stored RBCs. (B) Volcano plots of tests for storage differences for proteins, peptides, and PTMs. Select significant proteins, peptides, and PTMs are highlighted. Vertical black dashed lines at origin included for reference. Horizontal red dotted lines represent FDR < 5% threshold. (C) Examples of proteins with abundance differences between fresh and stored RBCs. SERPINF2 (left) is more abundant in fresh RBCs and PSMC6 (right) is more abundant in stored RBCs. Diagonal red dashed lines included for reference. (D) Gene ontology sets enriched in proteins that are significantly (FDR < 5%) more abundant in fresh RBCs (left) and stored RBCs (right). Top seven ontologies are shown for each. Vertical red dashed line represents FDR < 10% threshold for enrichment. (E) Proportions of PTMs that were significantly (FDR < 5%) more abundant in fresh or stored RBCs, stratified by modification type (*e.g.*, oxidation). Bracketed numbers represent p-values from hypergeometric tests for enrichment in fresh or stored RBCs. Red text highlights p-values < 0.05. (F) Comparison of the log10 ratio of PTMs with their unmodified peptides between stored and fresh RBCs. Select highly significant proteins are highlighted. Horizontal and vertical red dashed lines at the origin are included for reference. Diagonal gray dashed line included for reference. (G) Examples of proteins that possess a consistent ratios of PTM to its unmodified peptide in fresh and stored RBCs. An HBA peptide is more abundant in an unmodified state than a phosphorylated state (left) and an ALB peptide is more abundant in a glutathionylated state (right) in both fresh and stored RBCs. Diagonal red dashed line included for reference. (H) Volcano plots of tests for sex differences for proteins, peptides, and PTMs. Select significant proteins and peptides are highlighted. Vertical black dashed lines at origin included for reference. Horizontal red dotted lines represent FDR < 10% threshold. Horizontal black dotted lines represent lenient threshold of p-value < 0.05. (I) Comparison of sex coefficients between stored and fresh RBCs. Diagonal gray dashed lines included for reference. Black lines represent best fit regression lines comparing coefficients from stored and fresh RBCs. Proteins and peptides with significant sex differences (FDR < 10%) in both fresh and stored RBCs are highlighted with + symbols. (J) Gene ontology sets enriched in proteins that are significantly (FDR < 10%) more abundant in female (left) and male (right) RBCs, stratified by storage status. No gene ontologies were enriched for proteins more abundant in female fresh RBCs. Top seven ontologies are shown for each. Vertical red dashed line represents FDR < 10% threshold for enrichment.

### Storage alters proteolysis in RBCs

Storage lesion, a gradual degradation of RBCs during hypothermic storage, reduces their function and post-transfusion circulatory lifespan^9^. We identified 670 storage-responsive proteins (FDR < 0.05), with 397 elevated in fresh samples and 273 elevated in stored samples (**Figure 1B**). Gene set enrichment analysis (GSEA)^22^ found fresh samples to be enriched for immune-related proteins and negative regulators of proteolysis (*e.g.*, serpins^23^ – **Figure 1C**), while stored samples were enriched for proteases and proteasome components (**Figure 1D**). These findings indicate a significant role for protein degradation in the storage lesion^24^.

Changes in peptide abundance were largely consistent with changes at the protein level, highlighting differences in complement components (C3) and multiple apoplipoproteins (APO) involved in vesiculation events (a phenomenon exacerbated exponentially as storage progresses^9^), including APOA1, APOC3, APOE, and APOJ (clusterin or CLU) – a biomarker of RBC senescence and aging^25^ (**Figure 1B**). Comparing PTMs between fresh and stored RBCs (**Figure 1B, E**), we observed that oxidation and deamidation were the most abundant PTMs, and along with glutathionylation, differed significantly in frequency between fresh and stored RBCs. Of note, we observed an overall strong correlation between the levels of unmodified peptides with their matching PTMs (**Figure 1F**); examples include hemoglobin A1 peptides (HBA1 - mostly unmodified in fresh and stored RBCs) and albumin (ALB - mostly modified in fresh and stored RBCs) (**Figure 1G**). Notably, overall storage-associated changes in PTMs were higher magnitude and had higher significance than changes at the protein and (unmodified) peptide level.

Recent studies have highlighted a role for sex dimorphism in the RBC storage lesion^13^. Here we found few proteins with significant sex differences in abundance. Twelve proteins showed significant differences in fresh RBCs and a largely overlapping set of 10 in stored RBCs (**Figure 1H**). Sex-specific proteins were enriched for complement activation (**Figure 1J**), *e.g.*, elevated C9 in males, consistent with previously reported sex-specific differences across multiple tissues^26,27^, including the recent association of complement and coagulation components to stored RBC susceptibility to hemolysis^18^. While storage differences were most notable at the PTM level, sex differences were mostly observed at the protein and (unmodified) peptide level. Overall, the effects of storage and sex on proteins, peptides, and PTMs appeared independent.

### Genetic mapping of the RBC proteome

To identify genetic variants that influence protein traits, we applied genetic mapping analysis to proteins, unmodified peptides, and PTMs (**Figure 2A, B**; **Table S1; Supplementary Data File 1**). We mapped 954 protein QTL (pQTL; 165 *cis* and 764 *trans*) and 1,002 pQTL (197 *cis* and 787 *trans*) in fresh and stored RBCs, respectively. For peptides, we mapped 4,312 pepQTL (652 *cis* and 3,618 *trans*) and 4,847 (756 *cis* and 4,061 *trans*), representing 1,058 and 1,123 proteins in fresh and stored RBCs, respectively. For PTMs (after adjusting for their unmodified peptide levels), we mapped 668 ptmQTL (75 *cis* and 585 *trans*) and 731 (81 *cis* and 645 *trans*), representing 227 and 242 proteins in fresh and stored RBCs, respectively.

**Figure 2.**
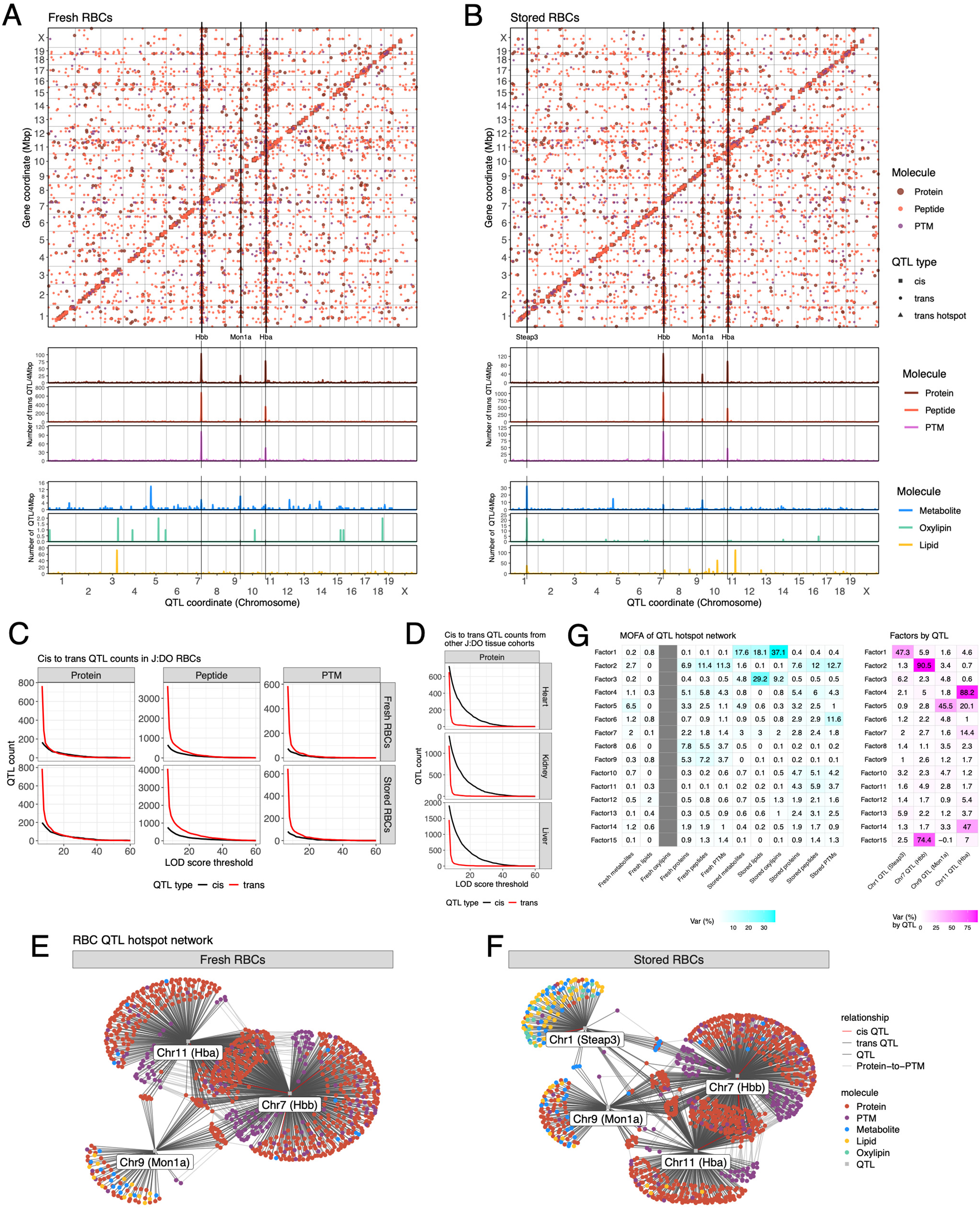
Genetic dissection of murine RBC proteome. QTL maps (LOD score > 7) for proteins, peptides, and PTMs in (A) fresh and (B) stored RBCs, aligned with QTL densities (based on a 4 Mbp window) for proteins, peptides, PTMs, metabolites, oxylipins, and lipids. Shape of data point indicates type of QTL (*e.g.*, *cis, trans,* hotspot). Vertical lines included to highlight QTL hotspots at the *Steap3*, *Hbb*, *Mon1a*, and *Hba* loci. More stringent QTL maps (LOD score > 8.3) are available in Figure S6B, C. (C) Comparison of detected QTL count by LOD score threshold for proteins, peptides, and PTMs in fresh and stored RBCs. Black lines represent *cis* QTL and red lines represent *trans* QTL. (D) Comparison of detected QTL count by LOD score threshold for proteins from heart, kidney, and liver tissues from two other J:DO cohorts. Black lines represent *cis* QTL and red lines represent *trans* QTL. QTL hotspot networks defined by three and four loci in (E) fresh and (F) stored RBCs, respectively. Color of node represents molecule type (*e.g.*, protein). Edge color represents relationships (*e.g.*, QTL type). (G) Heatmaps of the proportion of variance from multi-omics factor analysis (MOFA) for omic traits mapping QTL to the four hotspots in fresh and stored RBCs. Proportion of variance explained for factors by data layer (left) and by hotspot locus (right). The fresh oxylipins column is gray because no hotspot QTL were mapped.

One of the most striking features of RBC protein genetic architecture is the small number of *cis* QTL and relative enrichment of *trans* QTL compared to other tissues from J:DO mice^28–30^ (**Figure 2C, D**). In RBCs, *trans* QTL were 5 to 7-fold more prevalent than *cis* QTL. In contrast, other tissues had 5 to 6- fold more *cis* pQTL than *trans* pQTL. We observed stronger statistical associations (LOD scores) at *trans* QTL in RBCs, whereas in most tissues, *trans* associations tend to be weaker^31^. While the J:DO RBC cohort is approximately twice the size of the other tissue cohorts, which could contribute to the increased detection of *trans* QTL, it makes the reduction of *cis* QTL even more notable. Furthermore, *cis* QTL numbers are likely biased upward due peptides that contain coding variants^32^, which does not impact *trans* QTL (**Figure S1**). The predominance of *trans* effects is likely a signature of RBCs, which are denucleated and lack *de novo* protein synthesis^33^.

### QTL hotspots highlight key genes for RBC function and metabolism

The RBC *trans* pQTL were concentrated in four distinct hotspots, of which three were found in both fresh and stored RBCs (chromosome 7:120 Mbp, chromosome 9:107 Mbp, chromosome 11:33 Mbp) and one was unique to stored RBCs (chromosome 1:119 Mbp) (**Figure 2A, B, Figure S2**). Key genes involved in RBC function and metabolism are encoded in these hotspots, including *Steap3* (chromosome 1), hemoglobin beta (*Hbb* cluster, chromosome 7), *Mon1a* (chromosome 9), and hemoglobin alpha (*Hba* cluster, chromosome 11).

We aligned QTL results from our previous profiling of metabolites, oxylipins, and lipids from fresh and stored RBCs of this J:DO cohort^12,16,17^ (**Figure 2A, B, Figure S3A**). Metabolite, oxylipin, and lipid QTL mapped to the *Steap3* locus in stored RBCs. This was mirrored in higher heritability estimates in stored RBCs, most notably in metabolites and oxylipins (**Figure S3B**). Metabolite QTL also mapped to the *Hbb* and *Mon1a* hotspots. Many of these traits mapped to more than one hotspot (**Figure 2E, F**). From a biological standpoint, small molecules in this group mostly involve lipid and oxidized lipids that are enriched in RBC-derived vesicles, and guanine purine metabolites, involved in the regulation of GTPase-dependent vesiculation processes^34^. We performed multi-omics factor analysis (MOFA)^35^ on the combined molecular features that mapped to hotspots (**Figure 2G**). Several of the composite factors, representing the main axes of variation in fresh and stored RBCs, tracked closely with a QTL hotspot, further emphasizing the impact of these hotspots across multiple layers of RBC omic data. The impact of the hotspots on overall variation is still apparent when all features for fresh or stored RBCs are included (**Figure S4**).

### *Steap3* regulates apolipoproteins as part of its role in storage lesion moderation and ferroptosis

A gold standard to determine the quality of stored RBCs is post-transfusion recovery (PTR), a parameter that indicates the percentage of stored RBCs that still circulate at 24h after transfusion. Here we calculated PTR by determing survival of unlabeled stored RBCs in circulation, upon transfusion into Ubi-GFP RBC mice, and co-transfusion with tracer mCherry-labeled fresh RBCs. We previously identified *Steap3*, a ferrireductase, as the primary driver of PTR in stored murine RBCs^11^, including in J:DO mice^12^. High *Steap3* expressing alleles were associated with higher ferric iron reduction capacity, thus promoting iron loading during erythropoiesis^36–39^ and fueling Fenton chemistry in mature, stored RBCs by recycling ferric iron to its ferrous form upon oxidation. Here, we confirmed that coding variants in the *Steap3* gene are associated with altered STEAP3 protein expression by mapping a STEAP3 *cis* pepQTL (**Figure 3A, B**). The co-located PTR QTL was negatively correlated with STEAP3 expression, consistent with its hypothesized role in regulating storage lesion (**Figure 3C**), and the haplotype effects at the PTR QTL are negatively correlated with the STEAP3 pepQTL haplotype effects (**Figure 3D**).

**Figure 3.**
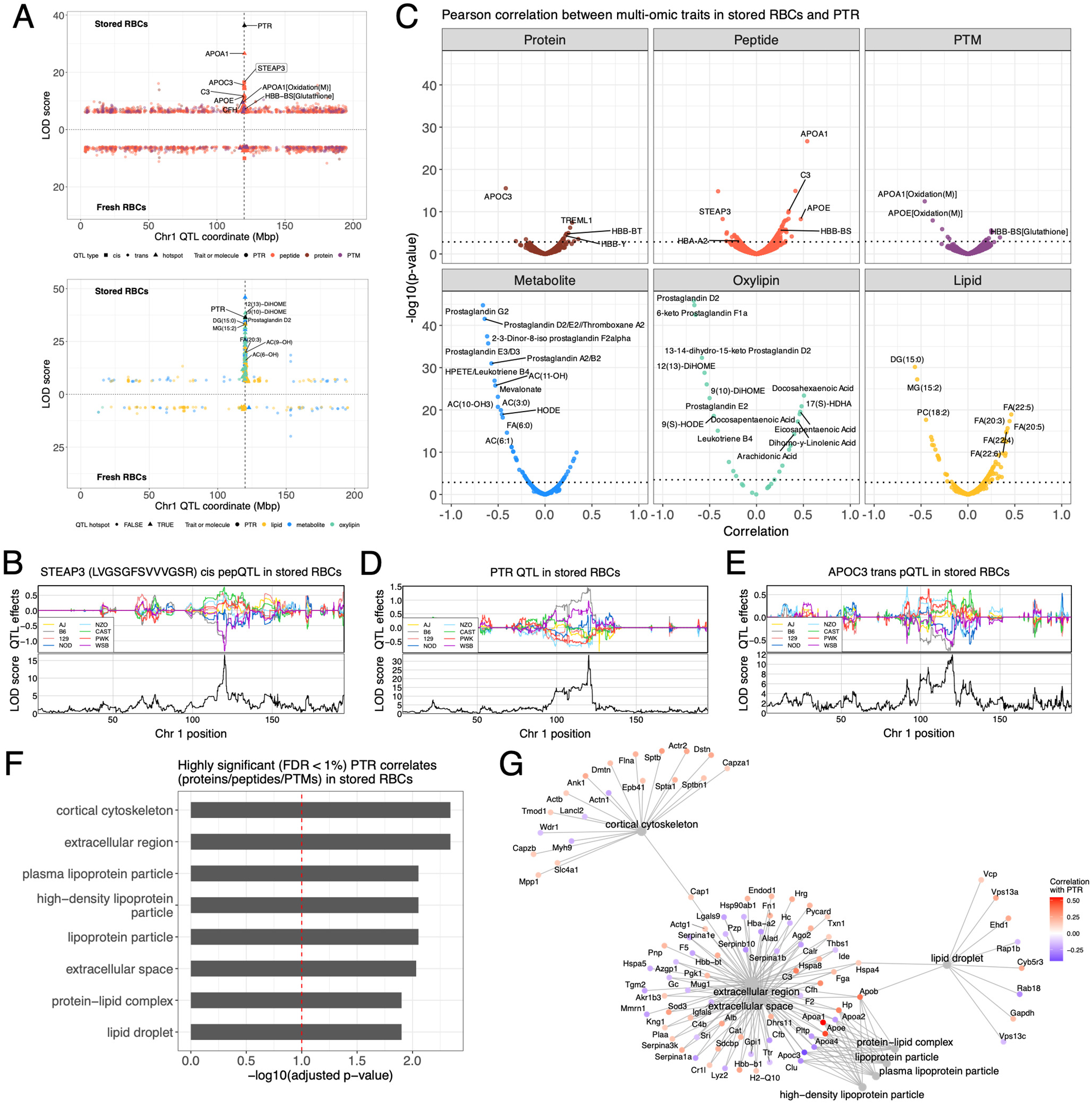
RBC proteomics data confirms STEAP3 as a driver of post-transfusion recovery (PTR) and implicates alipoproteins. (A) QTL hotspot maps to the *Steap3* locus on chromosome 1 and is specific to stored RBCs, comprising proteins, peptides, and PTMs (top) and metabolites, oxylipins, and lipids (bottom). (B) Haplotype effects for STEAP3 *cis* pepQTL. Association scores (LOD score) are aligned below the effects. (C) Volcano plots based on correlation t-test between omic traits and PTR. Select highly significant omic traits are highlighted. Horizontal red dotted lines represent FDR < 5% threshold. (D) Haplotype effects for PTR QTL that maps to the *Steap3* locus. Association scores (LOD score) are aligned below the effects. (E) Haplotype effects for apolipoprotein APOC3 *trans* pQTL that maps to the *Steap3* locus. Association scores (LOD score) are aligned below the effects. (F) Gene ontology sets enriched in proteins that are highly significantly correlated with PTR (FDR < 1%). Top eight sets are shown. Vertical red dashed line represents FDR < 10% threshold for enrichment. (G) Network plot of PTR-correlated proteins that belong to the top eight enriched gene sets. Color of gene node indicates the direction and magnitude of its correlation with PTR.

The *trans* QTL hotspot at the *Steap3* locus is seen only in the stored RBCs and it includes proteins involved in vesiculation processes (*e.g.*, several APOs), peptides, and redox PTMs, as well as metabolites, lipids, and oxylipins as previously reported^12^ (**Figure 3A**). The hotspot locus includes several apolipoproteins with haplotype effects that align with PTR dynamics: PTR was negatively correlated with inflammatory APOA2^40^ and APOC3^41^ (**Figure 3E**) and positively correlated with anti-inflammatory APOE^42^ and APOA1^43^. GSEA of protein correlates to PTR identified an enrichment for proteins involved in vesiculation events (**Figure 3F**), including proteins involved in actin cytoskeleton remodeling, lipid droplet formation and vesicle release in the extracellular space (**Figure 3G**). Of note, *Steap3* was first identified as a p53-dependent regulator of vesiculation in cancer^44–46^, suggesting a link between iron-dependent lipid and protein oxidation processes, induction of vesiculation in stored RBCs as drivers of the membrane-cytoskeletal remodeling leading to the morphological changes of stored RBCs^47^ from discocyte to echinocyte.

### Hemoglobin and *Mon1a trans* regulate proteasome components

Both hemoglobin alpha and beta chains map pQTL, pepQTL, and ptmQTL – especially oxidized and glutathionylated peptides, to the chromosome 1 hotspot, linking *Steap3* to the hemoglobin QTL hotspots on chromosomes 7 and 11 (**Figure 4A**). The *trans* QTL hotspots at *Hbb*, *Mon1a*, and *Hba* are present in both fresh and stored RBCs (**Figure 4A-C**). The extensive *trans* regulation observed through *Hbb* and *Hba* aligns with previous studies of hemoglobin-protein interactions^48,49^, highlighting cytosolic oxidation regulators, protein homeostasis, and metabolic enzymes. Most notably, several structural proteins mapped on the chromosome 7 *Hbb* hotspot, including Band 3 (SLC4A1, the most abundant RBC membrane protein that acts as a lynchpin of the RBC cytoskeleton to the membrane) and spectrin (SPTA1). On the other hand, top proteins mapping on chromosome 11 *Hba* included several antioxidant enzymes, like catalase (CAT), peroxiredoxin 6 (PRDX6), and glutathione peroxidase 1 (GPX1 – **Figure 4A, C**). Metabolites mapping to *Hbb* include glutathione disulfide and *S*-glutathionylcysteine, while IMP, GMP, GDP, and GTP map to *Mon1a* (**Figure 4B**).

**Figure 4.**
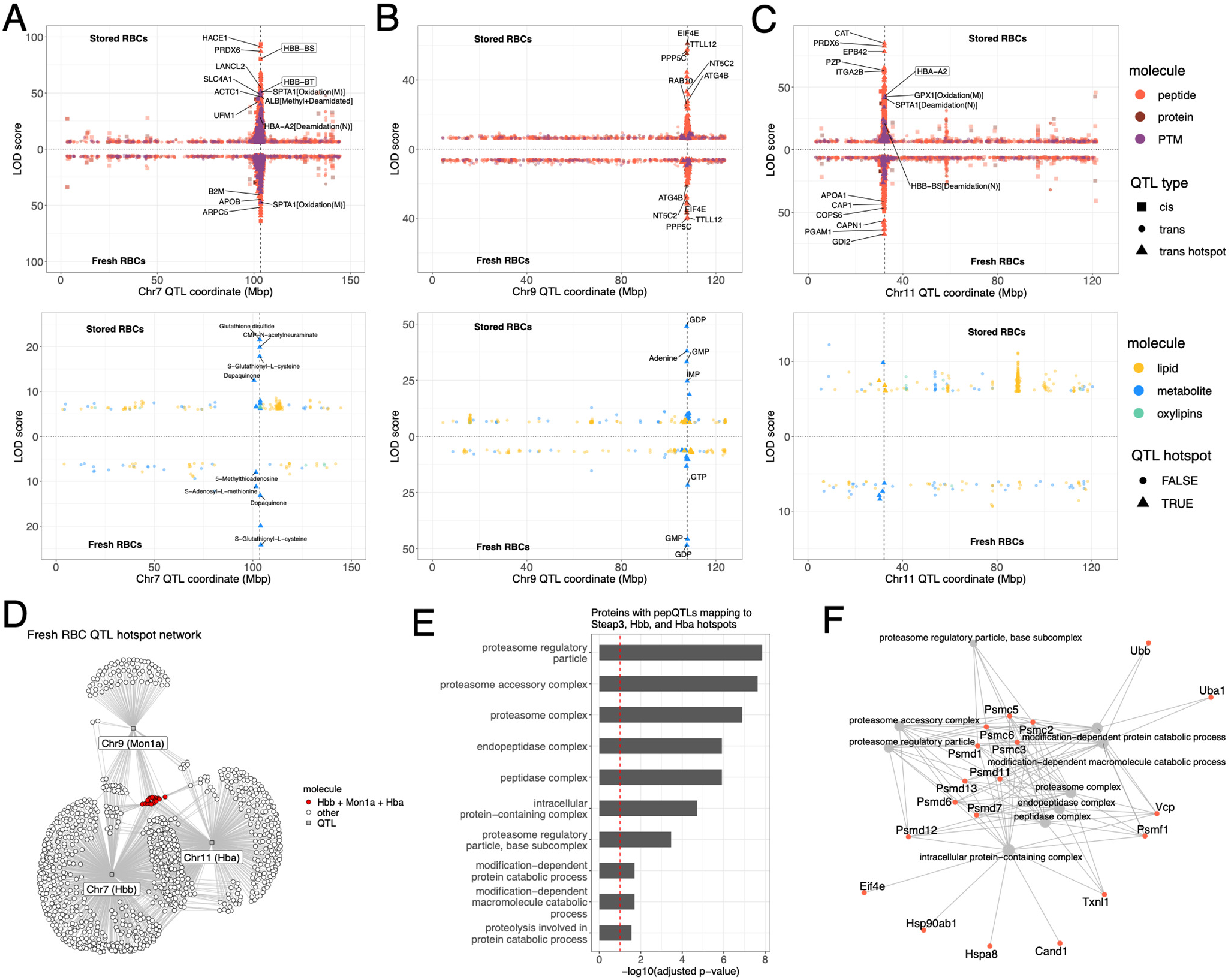
Proteomics and metabolomics reveal hemoglobin alpha and beta chains and *Mon1a* as connected regulatory hubs for RBC metabolism. QTL hotspots map to the (A) *Hbb* locus on chromosome 7, the (B) *Mon1a* locus on chromosome 9, and the (C) *Hba* locus on chromosome 9 in fresh and stored RBCs. Hotspots comprise proteins, peptides, and PTMs (top) and metabolites, oxylipins, and lipids (bottom). (D) QTL mapping networks defined by *Hbb*, *Mon1a*, and *Hba* loci in fresh RBCs. Proteins that map QTL to all three hotspot loci are highlighted as red. (E) Gene ontology sets enriched in proteins that map pepQTL to the three hotspots in fresh RBCs. Top ten sets are shown. Vertical red dashed line represents FDR < 10% threshold for enrichment. (F) Network plot of proteins with pepQTL that map to the three hotspots in fresh RBCs and belong to the top eight enriched gene sets.

*Mon1a* is functionally linked to hemoglobins through its role in vesicular trafficking of proteins involved in iron recycling from RBCs by spleen macrophages^50^. In the context of RBC storage, extravascular hemolysis^51^ via splenic erythrophagocytosis of morphologically altered stored RBCs^52^ is the main factor contributing to PTR. The 18 proteins that mapped pepQTL to all three hotspots (**Figure 4D**) are enriched in proteasome components and other protein catabolic proteins (**Figure 4E, F**), reflecting the importance of protein degradation and recycling in mitigating oxidative stress in RBCs and MON1A’s role in macrophage erythrophagocytosis.

### *Hbb* QTL hotspot driven by a novel cysteine variant

Mice carry two genes encoding hemoglobin beta chains (*Hbb-bt* and *Hbb-bs*)^53,54^. Some of the J:DO founder strains (AJ, 129, CAST, PWK, and WSB) possess only one cysteine residue (Cys93) in *Hbb-bt,* similar to the human HBB gene, while others (B6, NOD, and NZO), possess a second cysteine (Cys13, rs50836569) (**Figure 5A**). The haplotype effects of all the *trans* QTL that map to the *Hbb* hotspot, for example RHAG (**Figure 5B**), reflect the strain distribution of the *Hbb-bs* Cys13 allele among J:DO founder strains (**Figure 5C**).

**Figure 5.**
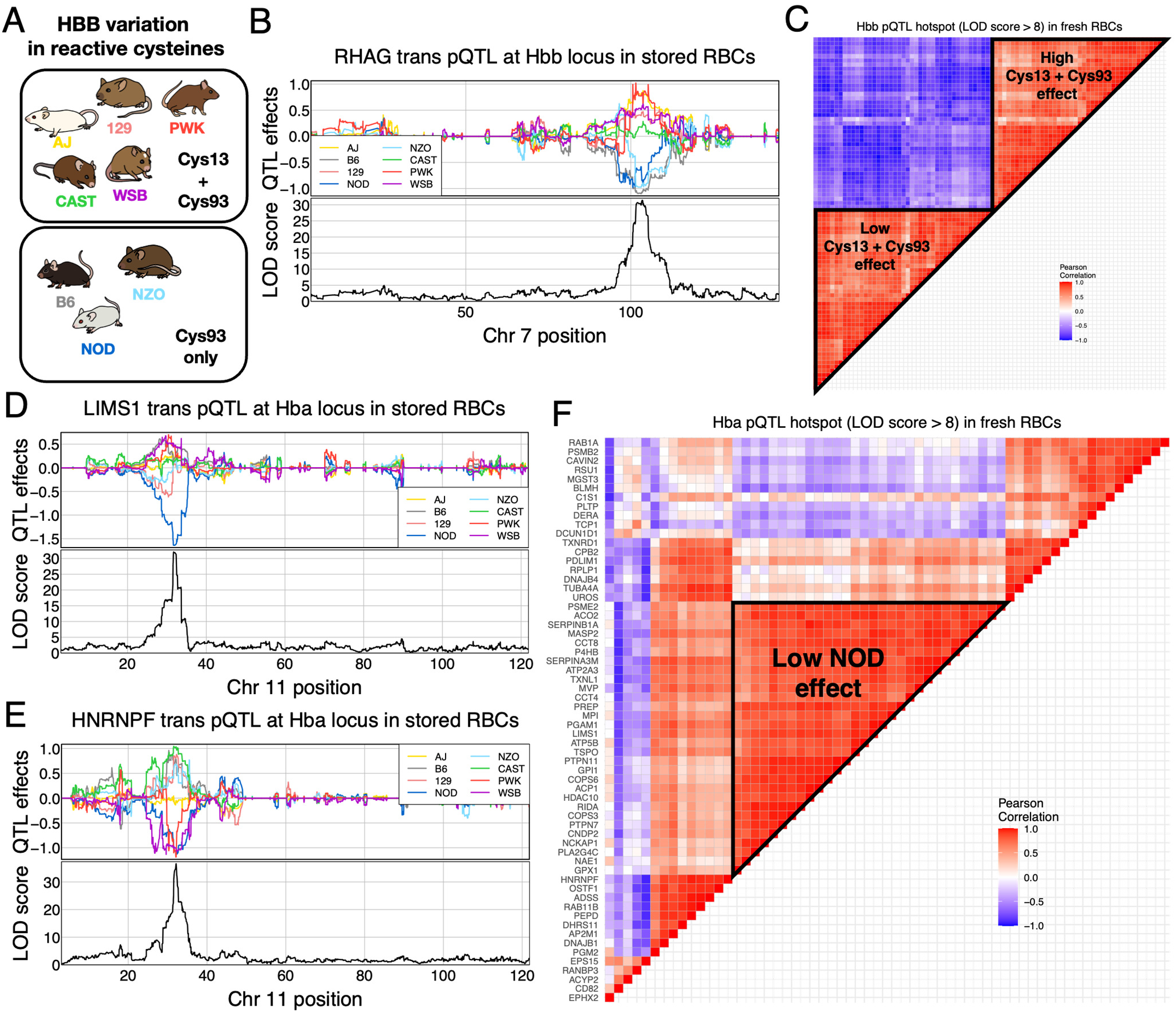
Genetic effects at hemoglobin alpha and beta loci. (A) Distribution of the Cys13 allele of *Hbb-bs* across the J:DO founder strains. (B) Haplotype effects for RHAG *trans* pQTL that maps to the *Hbb* locus. Association scores (LOD score) are aligned below the effects. (C) Heatmap of the pairwise correlations between haplotype effects for stringently detected pQTL (LOD score > 8) mapping to the *Hbb* locus. (D) Haplotype effects for LIMS1 *trans* pQTL that maps to the *Hba* locus. Association scores (LOD score) are aligned below the effects. (E) Haplotype effects for HNRNPF *trans* pQTL that maps to the *Hba* locus. Association scores (LOD score) are aligned below the effects. (F) Heatmap of the pairwise correlations between haplotype effects for stringently detected pQTL (LOD score > 8) mapping to the *Hba* locus. Triangle included to highlight the most prevalent effects pattern, characterized by a low NOD effect.

The haplotype effects of *trans* QTL that map to the Hba hotspot are more variable (**Figure 5F**), indicating that multiple genetic variants across the *Hba* gene cluster drive this hotspot. The most common *trans* pQTL haplotype effect pattern had a low NOD effect, as seen with LIMS1 (**Figure 5D**). Other *trans* pQTL show distinct haplotype effects, such as for HNRNPF (**Figure 5E**). Given this complexity and the annotated genetic variants in *Hba*, we could not determine obvious candidate variants for the *Hba* hotspot QTL. Further experimental follow-up would be needed to fine-map the causal genetic variants at *Hba*.

### Hemoglobin alpha and beta chains act as primary PTM hubs in RBCs

Stratifying ptmQTL by PTM group revealed oxidation and deamidation as the primary PTM groups that map *trans* ptmQTL to the *Hbb* and *Hba* hotspots (vertical bands on chromosomes 7 and 11 in **Figure 6A-B**), linking variation in hemoglobin chains to PTM variation in interacting proteins. The *Mon1a* hotspot maps fewer ptmQTL, suggesting different mechanisms compared to hemoglobin.

**Figure 6.**
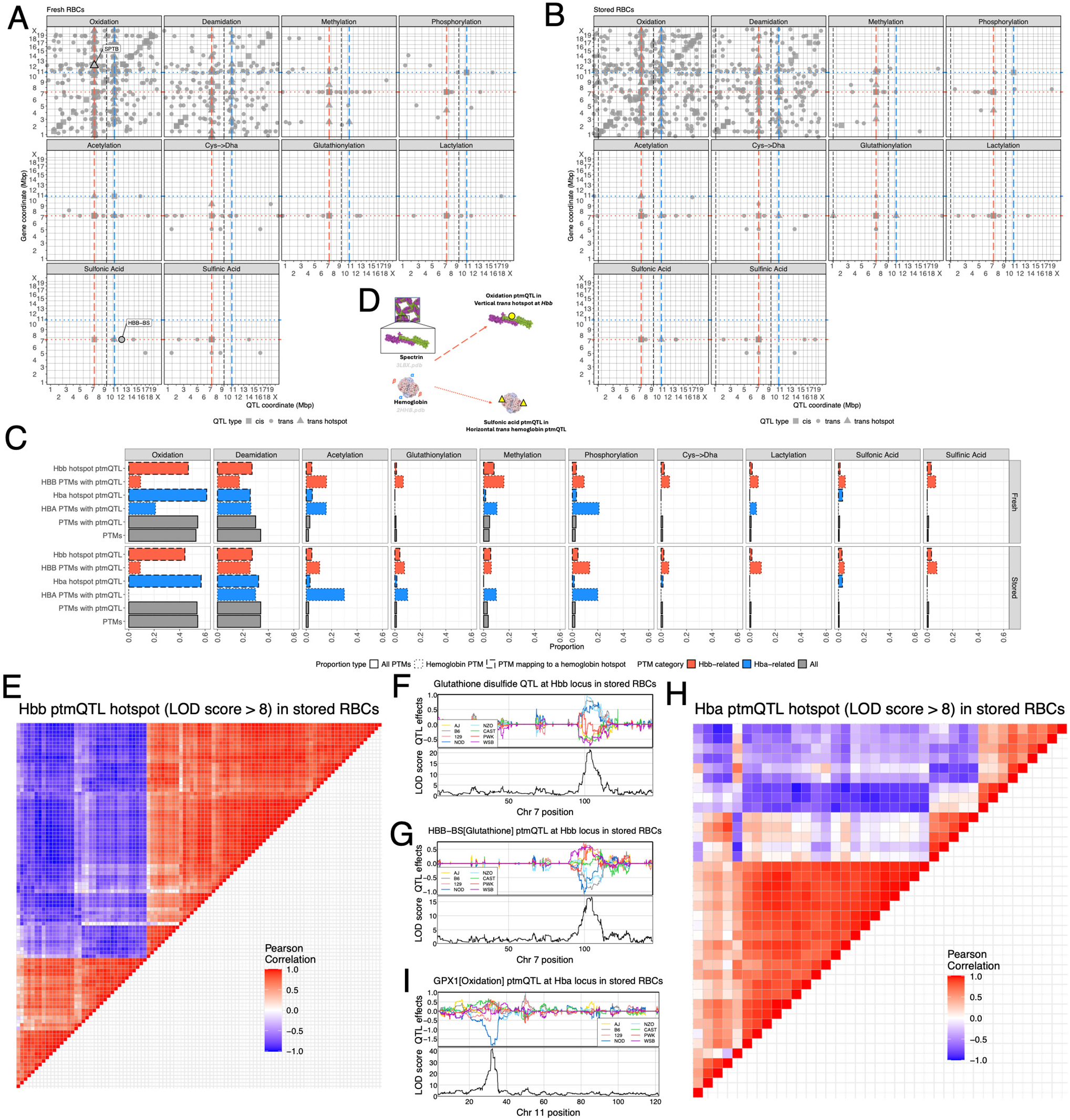
Hemoglobin alpha and beta chains are clear PTM hubs in RBCs. Maps of ptmQTL, stratified by PTM group, in (A) fresh and (B) stored RBCs. Shape of data point indicates type of QTL (e.g., cis, trans, hotspot). Vertical dashed lines represent ptmQTL hotspots at the Steap3, Hbb, Mon1a, and Hba loci. The Hbb and Hba loci are highlighted, colored red and blue, respectively. Horizontal dotted lines represents the PTMs of HBB (red) and HBA (blue). Example oxidation and sulfonic acid ptmQTL highlighted with labels, associated with Figure 6D. (C) Proportions of PTMs and ptmQTL, in reference to all PTM groups (gray), filtered to HBA PTMs and the Hba locus (blue), or filtered to HBB PTMs and the Hbb locus. Line type indicates whether a proportion is in reference to all PTMs (solid), those related to the Hbb or Hba ptmQTL hotspots (dashed), or those that are PTMs of HBB and HBA (dotted). (D) Diagram of potential interaction between hemoglobin and SPTB that results in oxidation of HBB and dioxidation (sulfonic acid) of SPTB. (E) Heatmap of the pairwise correlations between haplotype effects for stringently detected ptmQTL (LOD score > 8) mapping to the Hbb locus. (F) Haplotype effects for glutathione disulfide QTL that maps to the Hbb locus. Association scores (LOD score) are aligned below the effects. (G) Haplotype effects for a glutathione-bound HBB-BS peptide cis ptmQTL. Association scores (LOD score) are aligned below the effects. (H) Heatmap of the pairwise correlations between haplotype effects for stringently detected ptmQTL (LOD score > 8) mapping to the Hba locus. (I) Haplotype effects for an oxidized GPX1 peptide trans ptmQTL that maps to the Hba locus. Association scores (LOD score) are aligned below the effects.

The hemoglobin alpha and beta chain peptides also account for a high proportion of the ptmQTL that are not oxidation or deamidation (horizontal bands on chromosomes 7 and 11 in **Figures 6A-B** and **Figure 6C**), as well as ALB, encoded on chromosome 5. This highlights the wide-ranging effects of hemoglobin variation in RBCs, reflecting its central role in RBC function and metabolism. This dynamic between the *trans* ptmQTL hotspots at *Hbb* and *Hba* and hemoglobin peptide ptmQTL is illustrated with SPTB (**Figure 6D**), the spectrin beta chain, which is notably involved in stabilizing RBC membranes and implicated in oxidative modifications related to storage^55,56^. The *Hbb* ptmQTL hotspot allele effects align with the Cys13 variant (**Figure 6E**). We mapped a QTL for glutathione disulfide (GSSG), which had higher levels in mice possessing a single reactive cysteine residue in *Hbb-bs* (**Figure 6F**). A glutathionylated HBB-BS peptide mapped a *cis* ptmQTL with haplotype effects flipped relative to GSSG (**Figure 6G**), suggesting the extra cysteine potentially acts as a glutathione sink. The *Hba* hotspot, as above, shows QTL effect heterogeneity (**Figure 6H**). The *Hba* locus includes a *trans* ptmQTL for an oxidized GPX1 peptide (**Figure 6I**), which protects hemoglobin from oxidative stress^57^. These results highlight the extent to which hemoglobin interacts with other proteins and likely in regulating PTMs.

### Humanized *Hbb* mouse model

To study the impacts of *Hbb* on the RBC proteome and metabolome, we created genetically modified mice with altered *Hbb* cysteines. In humans, Cys93 helps buffer intracellular glutathione during hypoxia or oxidative stress^58^ and facilitates the recycling of oxidized peroxiredoxin-2 dimers^59^. It may also regulate vasodilation by binding nitric oxide^60,61^.

We investigated the metabolome, proteome, and redox PTMs (including protein Cys-glutathionylation) in three groups (with three mice each): B6 (WT) with one cysteine at position 93 for HBB-BS, humanized mice expressing human canonical HBB (with Cys93 only) and no murine HBB-BS or HBB-BT, and humanized mice expressing HBB C93A (Cys93Ala) and no murine HBB-BS or HBB-BT (**Figure 7A**). After confirming the absence of murine HBB-BS or HBB-BT and the exclusive expression of human HBB in the humanized mice (**Figure 7B, C**), we evaluated PTR. Both humanized strains showed a significant drop in PTR (**Figure 7D**), but only the C93A mice lower intracellular levels of reduced glutathione (GSH) (**Figure 7E**).

**Figure 7.**
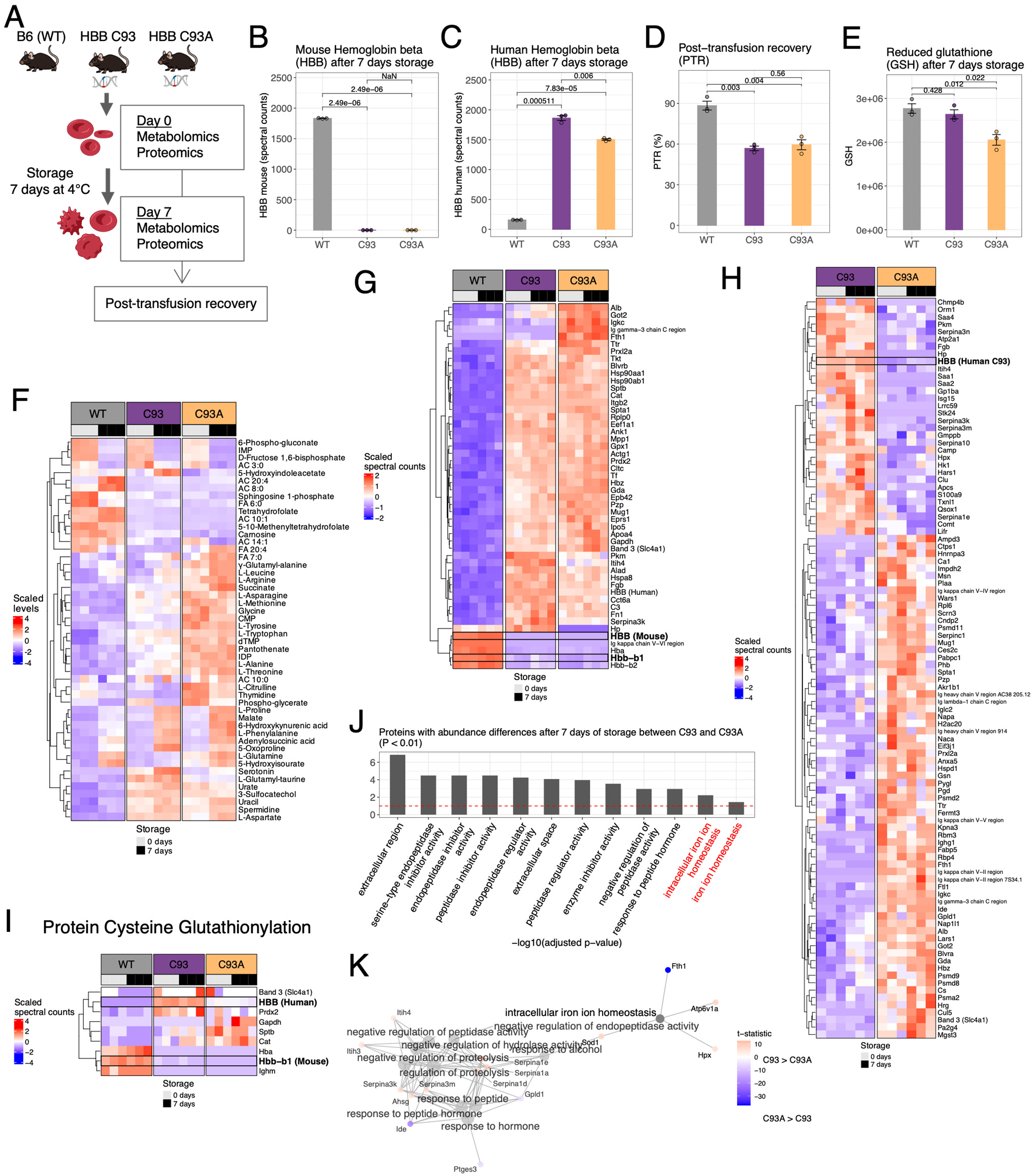
Loss of cysteine residue in humanized *Hbb* mouse model impact proteins related to proteolysis and iron homeostasis. (A) Diagram of the humanized *Hbb* mouse model experiment, including mouse strains and phenotypes measured. The mouse cohort is composed of three mice each from the reference B6 strain (WT), humanized *Hbb* mice (C93), and humanized *Hbb* mice with the C93A mutation (C93A). Comparisons of (B) mouse HBB levels (as spectral counts) after seven days of storage, (C) of human HBB levels (as spectral counts) after seven days of storage, (D) post-transfusion recovery (PTR), and (E) reduced glutathionylation (as spectral counts) after seven days of storage across mice. Standard error bars on group means and p-values from pairwise t-tests included. (F) Heatmap of scaled metabolite levels across groups and storage times. (G) Heatmap of scaled protein levels across groups and storage times. Hbb-b1 and mouse HBB (Hbb-b1 + Hbb-b2) are highlighted. (H) Heatmap of scaled protein levels for humanized mouse groups and storage times. Human HBB is highlighted. (I) Heatmap of cysteine glutathionylation levels across groups and storage times. Human HBB and mouse Hbb-b1 are highlighted. (J) Gene ontology sets enriched in proteins with abundance differences between C93 and C93A mice after seven days of storage (p < 0.01). Top ten sets are shown, as well as two sets related to iron metabolism, highlighted with red text. Horizontal red dashed line represents FDR < 10% threshold for enrichment. (K) Network plot of proteins with abundance differences between C93 and C93A mice after seven days of storage. Color of gene node indicates the direction and magnitude of the t-statistic comparing group means. The intracellular iron ion homeostasis gene set is emphasized with greater opacity.

Metabolomics profiling showed that human HBB expression is associated with storage-related changes in glutathione homeostasis (accumulation of 5-oxoproline), purine oxidation (urate accumulation, even in fresh RBCs), and altered carboxylic acid (malate). These effects were more pronounced in C93A mice, which had higher levels of purine oxidation products (5-hydroxyisourate), carboxylic acids (succinate – derived from succinyl-CoA, precursor to heme synthesis), energy metabolism (phosphoglycerate isomers), and amino acids (alanine, arginine, asparagine, glycine, glutamine, leucine, methionine, threonine, tryptophan, tyrosine, citrulline), consistent with systemic alterations to protein synthesis/degradation (**Figure 7F**).

Proteomics analyses revealed widespread changes in the proteome of Hb-humanized mice compared to WT mice (**Figure 7G**). Direct comparison of the canonical and HBB C93A RBCs confirmed the lack of expression of the C93-containing peptide in the C93A mice, as well as widespread effects on the proteome (**Figure 7H**) and glutathionylated peptidome (**Figure 7I**) due to the loss of C93, consistent with the pQTL and ptmQTL results. Notably, HBB cysteine status constrained glutathionylation at critical functional residues, such as GAPDH C152 in the active site of this rate-limiting glycolytic enzyme^62^ and Band 3 (SLC4A1) at DIDS-sensitive C, which does not regulate anion homeostasis^63^. GSEA for proteins with differing expression between C93 and C93A mice after seven days of storage (P < 0.01; **Figure 7J- K**) revealed pathways related to proteins that change with RBC storage or that map QTL to all hotspots (peptidase activity and proteasome components) and the *Steap3* QTL hotspot (iron metabolism and homeostasis).

## DISCUSSION

Inconsistencies between transcriptional and proteomic data have challenged the central dogma of molecular biology^64^, shifting the focus from a one-gene one-protein model to the study of proteoforms^65^, which arise from variations in gene expression, post-transcriptional, and post-translational modifications. We used mass-spectrometry to quantify proteins, peptides, and their PTMs in RBCs, a unique cell type that lacks nuclei and organelles, and thus is incapable of *de novo* protein synthesis. Genetic mapping of proteome traits revealed that RBCs, unlike other tissues, exhibit strong *trans*-regulation of intracellular protein abundance, specifically at three hotspots that colocate with the genes *Hba*, *Hbb*, and *Mon1a*.

The J:DO population is a powerful resource for investigating the impact of genetic variance on RBC biology, revealing for the first time the system-wide impact of genetic variants at the hemoglobin loci on traits, linking gas transport, redox homeostasis, and proteostasis in mature erythrocytes. Genetic variation in the *Hbb* locus is central to RBC proteome homeostasis and contributes to the intensity of hemolysis observed in Mendelian hemolytic diseases, such as sickle cell disease or beta-thalassemia (E6V^66,67^). Over 3,105 HBB variants are listed in the DisGenet database^68^ (v24.2), including the HBB:c.281G>T missense mutation, which substitutes phenylalanine for cysteine in humans^67^, making the C93A experiments in mice immediately relevant. During oxidative stress, such as blood storage, beta-elimination of C93 thiol forms dehydroalanine, losing sulfur biochemistry and the capacity to promote recycling of peroxiredoxin dimers^59^. HBB irreversibly oxidizes peptides, including Cys → DHA mapped to Steap3. By constraining intracellular iron levels during erythropoiesis, STEAP3 regulates hemoglobin concentrations, with genetic ablation resulting in hypochromic anemia in mice and humans^36–39^.

While polymorphisms in the region coding for *Steap3* were previously associated with poor storage quality and elevated lipid peroxidation^11,12^, this is the first report linking genetic variation to STEAP3 protein levels, PTR, protein glutathionylation, and oxidation of RBC proteins, including hemoglobins. Pathway analysis of proteins correlated to PTR highlighted not just STEAP3, but also metabolic enzymes involved in glycolysis (HK1, PFKM, PKLR, BSG – essential for lactate transporter relocation to the membrane upon its synthesis^69^), the pentose phosphate pathway (RPIA, G6PDX, PGLS, PGD, TALDO1, PRPS1), ferroptosis (STEAP3, GPX1, GPX4, HBB-y, NQO1, PRDX2, GSR, SOD3, CAT), all consistent with recent literature in humans^12–17^. Novel findings here include the identification of extracellular or vesiculation-related proteins, cross-linking enzyme (TGM2^70^) complement and coagulation components (C3, F5, FGA, HP, APOE, APOA1, APOA2, APOA4)^18^, calcium signaling (CALR), heat shock proteins (HSP90AB1, HSPA5, HSPA8^71,72^) and proteasome (PSMC2,5,6; PSMD1,6,7,11,12,13; PSMF1)^24^ or ubiquitination components (VCP, UBA1, UBB, CAND1)^73^, all previously associated with increased hemolytic propensity in response to hypoxia or hematological disorders, like sickle cell disease, beta-thalassemia, or chorea acanthocytosis. These proteins all mapped QTL to the *Steap3, Hbb*, and *Hba* hotspots.

Overall, these results suggest that HBB levels and their cysteine residues are master regulators of intracellular redox homeostasis beyond serving as a simple buffer for RBC glutathione^53,58^. This view aligns with mathematical predictions of the total oxidative stress burden that circulating (or stored) RBCs may face, which is a function of hemoglobin autoxidation and directly proportional to the variance in the total number of hemoglobin molecules per cell. It follows that iron metabolism and mean cell hemoglobin concentration, as well as genetic or other factors regulating them, such as blood donation frequency or repletion^74^, could contribute to inter-donor heterogeneity in hemolytic propensity and RBC susceptibility to hemolysis in general, making our findings relevant beyond transfusion medicine.

Altogether, our results strengthen the connection between STEAP3 and HBB in RBC response to storage highlighting a complex feedback loop between these circuits, as mediated by: (i) a cross-talk between storage-associated iron metabolism (linked to *Steap3*); (ii) oxidant stress and redox damage to proteins and lipids (promoted by STEAP3 and counteracted by glutathione pools); (iii) availability of glutathione pools (constrained by HBB cysteine); (iv) HBB cysteine redox status (cross-regulated by STEAP3); (v) proteasomal degradation of oxidized components (QTL mapping to both *Hbb* and *Steap3*); and (vi) vesiculation of irreversibly damaged components (*Steap3,* APOs, *Mon1a* mapping guanosine phosphate pools).

This study provides a novel comprehensive RBC resource for viewing system-wide impacts of genetic perturbations in a genetically diverse population, offering a unique platform for future studies of RBCs relevant to the hematology community.

## STAR METHODS

### J:DO Mouse cohort

We acquired 600 J:DO^19^ mice from the Jackson Laboratory, representing the 45^th^ and 46^th^ generations of the population. The mice were shipped in three male-only batches and three female-only batches, each batch comprised of 100 animals. 350 animals were selected from the 1^st^ through 5^th^ batches for RBC sample collection. This subset included 202 males and 148 females. RBC samples were collected, and post-transfusion recovery (PTR) measured based on comparing RBCs under fresh and storage conditions for each individual. Fresh and stored RBC samples underwent mass spectrometry quantification of proteins and PTMs. Mass spectrometry for metabolites, oxylipins, and lipids were described previously^12,16,17^. One male mouse’s stored RBC sample was lost, resulting in 349 individuals for the stored data.

### RBC sample collection, storage, and PTR measurement

Murine RBC storage, transfusion, and PTR determinations were carried out as previously described^75^, modified as described in Hay et al. 2023^76^. Whole blood was drawn by cardiac puncture under sterile conditions into CPDA-1, was centrifuged, and the hematocrit was adjusted to 75% by removing supernatant. Blood was stored at 4°C for 7 days. RBCs from J:DO mice were used as the “test” population. Ubi-GFP mice were used as recipients to allow visualization of the test cells in the non-fluorescent gate. To control for differences in transfusion and phlebotomy, RBCs from ROSA26-LCB-mCHERRY mice (mCHERRY) were used as a tracer RBC population (never stored) was added to stored RBCs immediately prior to transfusion. PTR was calculated by dividing the post-transfusion ratio (Test/Tracer) by the pre-transfusion ratio (Test/Tracer^75^). At the time of transfusion, blood samples were frozen in liquid nitrogen and stored at −80°C until subsequent analysis. Humanized mice expressing the canonical human HBB (β93Cys) or HBB C93A (β93Ala) were obtained from Dr. Rakesh Patel (University of Alabama at Birmingham), and were previously described^59,77^. Data were collected from three animals per each group (B6 WT, canonical HBB, and HBB C93A). All experimental protocols were approved by the University of Virginia IACUC on 04/22/2019 (protocol n: 4269).

### Mass spectrometry for RBC proteomics

Proteomics analyses were performed as described previously^78^. A volume of 10 μL of RBCs were lysed in 90 μL of distilled water containing 10 mM *N*- ethylmaleimide (NEM) for 20 min at room termperature. 5 μL of lysed RBCs were mixed with 45 μL of 5% SDS and then vortexed. Samples were reduced with 10 mM DTT at 55 °C for 30 min, cooled to room temperature, and then alkylated with 25 mM iodoacetamide in the dark for 30 min. Next, a final concentration of 1.2% phosphoric acid and then six volumes of binding buffer (90% methanol; 100 mM triethylammonium bicarbonate, TEAB; pH 7.1) were added to each sample. After gentle mixing, the protein solution was loaded to a S-Trap 96-well plate, spun at 1500 x g for 2 min, and the flow-through collected and reloaded onto the 96-well plate. This step was repeated three times, and then the 96-well plate was washed with 200 μL of binding buffer 3 times. Finally, 1 μg of sequencing-grade trypsin (Promega) and 125 μL of digestion buffer (50 mM TEAB) were added onto the filter and digested carried out at 37 °C for 6 h. To elute peptides, three stepwise buffers were applied, with 100 μL of each with one more repeat, including 50 mM TEAB, 0.2% formic acid (FA), and 50% acetonitrile and 0.2% FA. The peptide solutions were pooled, lyophilized, and resuspended in 500 μL of 0.1 % FA. Each sample was loaded onto individual Evotips for desalting and then washed with 200 μL 0.1% FA followed by the addition of 100 μL storage solvent (0.1% FA) to keep the Evotips wet until analysis. The Evosep One system (Evosep, Odense, Denmark) was used to separate peptides on a Pepsep column, (150 um inter diameter, 15 cm) packed with ReproSil C18 1.9 µm, 120A resin. The system was coupled to a timsTOF Pro mass spectrometer (Bruker Daltonics, Bremen, Germany) via a nano-electrospray ion source (Captive Spray, Bruker Daltonics). The mass spectrometer was operated in PASEF mode. The ramp time was set to 100 ms and 10 PASEF MS/MS scans per topN acquisition cycle were acquired. MS and MS/MS spectra were recorded from m/z 100 to 1700. The ion mobility was scanned from 0.7 to 1.50 Vs/cm^2^. Precursors for data-dependent acquisition were isolated within ± 1 Th and fragmented with an ion mobility-dependent collision energy, which was linearly increased from 20 to 59 eV in positive mode. Low-abundance precursor ions with an intensity above a threshold of 500 counts but below a target value of 20000 counts were repeatedly scheduled and otherwise dynamically excluded for 0.4 min.

### Normalizing proteomics data

We applied a log_10_(*x* + 1) transformation to each trait (protein, peptide, or PTM). We evaluated whether to convert zeros in the data to missing values (NAs) by comparing heritability estimates, fit with the qtl2 R package^79^. Heritability represents the proportion of variability explained by genetic relatedness within a population. We observed consistently higher heritability for the data with missing values compared to zeros, suggesting that a large proportion of NAs/zeros are more consistent with missing at random rather than zeros or left censoring. As such, we treated them as missing for downstream analysis.

### Testing proteins, peptides, and PTMs for abundance differences between fresh and stored RBCs

Given that fresh and stored samples was collected for all individuals, we tested for proteins, peptides, and PTMs that differed in abundance between fresh or stored RBC status using a paired t-test. P-values were then false discovery rate (FDR)-adjusted using the Benjamini-Hochberg method^80^ within each data type (*e.g.*, proteins) separately.

### Testing whether a peptide was primarily present in a modified or unmodified state

We also sought to detect PTMs that systematically differed in abundance from their unmodified peptides within fresh and stored RBCs. We performed a paired t-test comparing each PTM’s abundance to its unmodified peptide. P-values were then FDR-adjusted using the Benjamini-Hochberg method^80^ within fresh and stored RBCs separately.

### PTM adjustment for peptide

We were primarily interested in sex differences and ptmQTL for PTMs that were distinct from their unmodified peptide (**Figure S5**). We used an regression approach to pre-adjust for peptide, similar to what we did previously for phosphorylation data^81^:

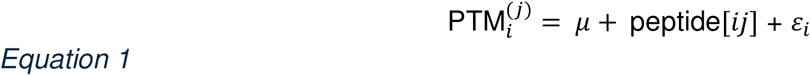

where 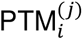 is the abundance level for PTM *j* for individual *i*, *μ* is the intercept, peptide[*ij*] is the contribution of the corresponding unmodified peptide for PTM *j* for individual *i*, and *ε_i_* is the error for individual *i*, modeled as ε*_i_* ∼ N(0, *σ*^2^). An adjusted PTM was then calculated as a residual: 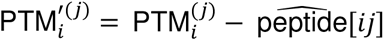, where 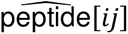 is a fixed effect prediction.

### Testing proteins, peptides, and PTMs for sex differences in abundance

To reduce the potential for false positive sex differences, we filtered out proteins, peptides, and PTMs with fewer than 100 non-missing observations. We also further transformed the traits, converting the ranks to normal quantiles (*i.e.*, rank-based inverse normal transformation) to conservatively reduce the influence of outliers.

One challenge for testing sex differences in this study’s J:DO cohort is that batch and sex are partially confounded because each batch is comprised of a single sex. This is an issue because batch effects often drive extensive variation in large-scale proteomics studies^82^. Nevertheless, consistency across sex-specific batches would provide statistical evidence of sex differences which can be assessed with a linear model. We fit a linear mixed effect model (LMM) with a random effect to account for batch:

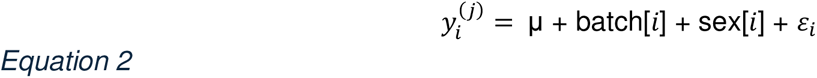

where 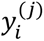 is the normal quantile for trait *j* for individual *i*, *μ* is the intercept, batch[*i*] is the contribution of the corresponding batch for individual *i*, sex[*i*] is the contribution of individual *i*’s sex, and ε_*i*_ is the error for individual *i*, modeled as ε_*i*_ ∼ N(0, *σ*^2^). Sex is modeled as a fixed effect coefficient, whereas batch is fit as a random effect: batch_*b*_∼ N(0, τ^2^). The LMM was fit using the lmer function from the lme4 R package^83^. The sex coefficient was then tested using ANOVA, comparing the model in Equation 2 to one excluding the sex covariate, with the lmerTest R package^84^ loaded. P-values were then FDR-adjusted using the Benjamini-Hochberg method^80^ within data type and fresh and stored RBCs separately.

### QTL analysis

As with testing for sex differences, we filtered to proteins, peptides, and PTMs with at least 100 non-missing observations and used the rank-based inverse normal transformation, conservatively converting all traits to standardized normal quantiles. To map QTL, we used the qtl2 R package^79^ which is commonly used for genetic mapping in multiparental populations (MPPs)^85^, such as the J:DO. For each trait we performed a QTL scan, in which the following LMM is fit at loci spanning the genome:

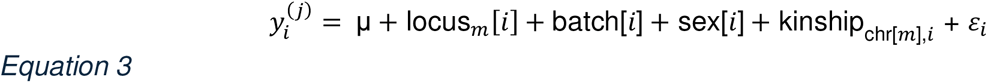

where 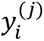 is the normal quantile for trait *j* for individual *i*, *μ* is the intercept, locus_*m*_[*i*] is the effects of genetic locus *m* on individual *i*, batch[*i*] and sex[*i*] are the contributions of batch and sex on individual *i*, fit as fixed effect covariates, kinship_chr[*m*],*i*_ and ε_*i*_ are random terms representing a polygenic effect that can account for population structure and unstructured error, respectively: kinship_chr[*m*],*i*_ ∼ N(**0**, **G**_*m*_*τ*^2^) and ε_*i*_ ∼ N(0, *σ*^2^). **G**_*m*_ is an *n* × *n* genetic relationship matrix estimated from all markers excluding those from the chromosome of locus *m*, commonly referred to as leave-one-chromosome-out (LOCO)^86,87^. LOCO increases QTL mapping power by restricting the random kinship term from absorbing signal from the tested locus term^88^. We also estimated narrow-sense heritability, the proportion of phenotypic variation explained by additive genetic effects using the qtl2 R package^79^. Briefly, heritability is estimated using a modified version of Equation 3, excluding the locus term and using a non-LOCO kinship matrix (*i.e.*, estimated from all markers). Heritability is than calculated as 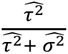.

### QTL model

The conventional QTL mapping approach in MPPs is to reconstruct the genotypes at a locus in terms of the founder haplotypes rather than a SNP (eight founder haplotypes versus two SNP alleles) using a hidden Markov model and the eight founder strains’ genotypes^79,89^. The locus term is modeled as 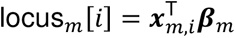 where 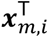 is 1 × 8 additive founder haplotype probability vector at locus *m* for individual *i* and **β**_*m*_ is the 8 × 1 founder haplotype effect vector at locus *m*. During genome scans, the locus terms are modeled as fixed effects for the sake of computational efficiency. However, when interpreting founder haplotype effects at QTL (*e.g.*, **Figure 3B**), we re-fit the locus term as a random effect and then calculate best linear unbiased estimates (BLUPs) of **β**_*m*_, which conservatively constrains potentially unstable fixed effect regression coefficients.

### QTL significance threshold

During QTL genome scan, each locus is tested by comparing the model in Equation 3 to one excluding the locus term using a logarithm of odds (LOD) score statistic, roughly analogous to a likelihood ratio. Giving specific thresholds and their error rates is impossible because they will vary with the sample population (*e.g.*, number of recombinations, varying polygenic background), we instead used a lenient LOD score threshold of 7 (genome-wide type I error of 0.36 based on simulations) to allow the detection of subtle QTL that were further supported by external biological information (*e.g.*, *cis* signal and *trans* hotspots). A LOD score thresholds of 8.3 corresponds to a genome-wide type I error of 0.05, which we found still resulted in the reported QTL hotspots (**Figure S6**). Genome-wide type I error rates were estimated from 1,000 null simulations (with background polygenic effect of 50%) from the genomes of the 350 J:DO mice used in this study using the musppr R package^90^.

Detected QTL were defined as *cis* if the encoded gene’s start position was within 10 Mbp of the peak QTL position. We use this wide *cis* window because the J:DO have greater linkage disequilibrium than natural populations. The genotypes and gene coordinates followed GRCm39.

### Multi-Omics Factor Analysis (MOFA)

We performed MOFA using the MOFA2 R package^35^. We first performed an iterative Combat normalization with the pcaMethods R package^91^ to adjust for batch effects in each data layer (metabolites, lipids, oxylipins, proteins, peptides, and PTMs) for both fresh and stored RBCs. We then use MOFA with data filtered to the traits that mapped QTL to the four hotspots to estimate 15 factors. For each factor, we calculated the proportion of variance explained by each of the hotspot loci. This process was repeated for the fresh and stored data, separately, to estimate 26 factors with no filtering to traits that mapped hotspot QTL.

### Gene set enrichment analysis

We performed GSEA using the clusterProfiler R package^22^. We defined gene sets (using ENSEMBL gene ID) based on proteins with storage differences, sex differences, or QTL. The stringency of the threshold to define sets varied depending on the extent of signal. The background universe was set to all genes with an observed protein, peptide, or PTM. All ontologies – biological process (BP), molecular function (MF), and cellular component (CC) – were used. An FDR-adjusted p-value threshold of 0.1 was used for determining significant ontologies.

### Testing for protein abundance differences between C93 and C93A humanized HBB mice

We tested for protein abundance differences between C93 and C93A mice using an unpaired t-test for fresh (0 days) and stored (7 days) samples.

### Software

All analysis was performed using the R statistical programming language^92^.

## Data availability

All data and R code to generate reported results are available at https://doi.org/10.6084/m9.figshare.28465025. Processed data and results are available at https://churchilllab.jax.org/qtlviewer/Zimring, where users can also perform interactive web-based analysis through the QTLViewer^93^. Proteomics data associated with this study are publicly available through the MassIVE repository (ID: MSV000095934).

## Acknowledgements

AD and JCZ were supported by funds by the National Heart, Lung, and Blood Institute (NHLBI) (R21HL150032, R01HL146442, R01HL149714, R01HL148151). G.R.K and G.A.C were supported by grants from the National Institute of General Medical Sciences (NIGMS), F32GM124599 and R01GM070683, respectively. The content is solely the responsibility of the authors and does not necessarily represent the official views of the National Institutes of Health. We thank Corinne M. Keele for her illustrations of the J:DO founder strains (**Figure 1.A**).

## Authorship Contribution

Animal studies: AH, JCZ. Proteomics: MD, KCH. Metabolomics analyses: TN, DS, JAR, AD. Biostatistics and Bioinformatics: GRK, CO’C, MV, GPP, GAC. pQTL analyses: GRK, GAC. Figure preparation: GRK, AD. Writing: GRK, GAC, AD.

## SUPPLEMENTARY FIGURES

**Figure S1.**
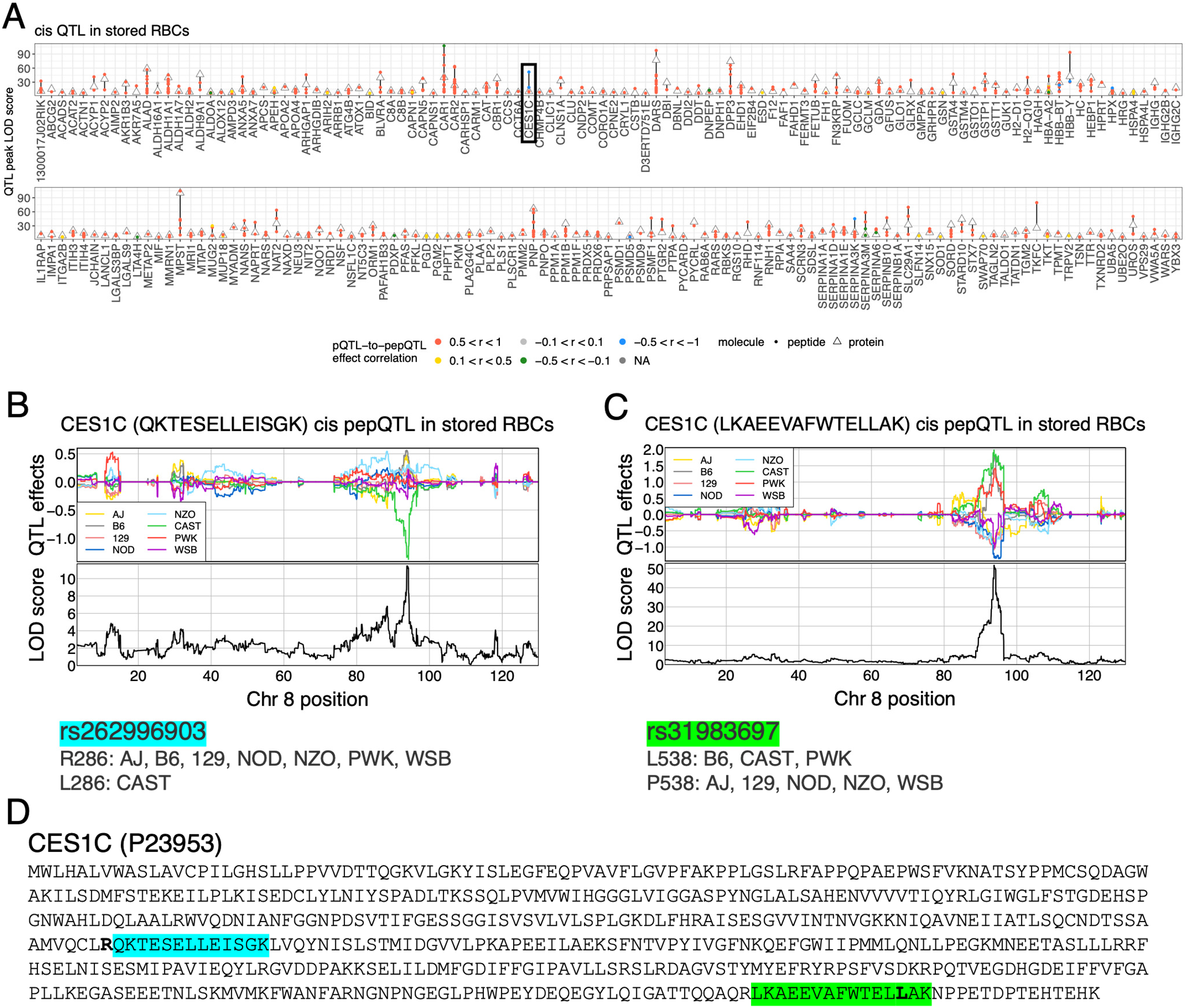
Coding variation can reduce power and drive false abundance pQTL and pepQTL. (A) LOD scores for proteins that map both *cis* pQTL and *cis* pepQTL in stored RBCs. Triangle points represent pQTL LOD scores. Circle points represent pepQTL LOD scores and are colored based on their haplotype effect correlation with the corresponding pQTL. CES1C is highlighted with black rectangle, which is explored in **Figure S1B-D**. (B) Haplotype effects for CES1C *cis* pepQTL for a peptide with QKTESELLEISGK sequence. Association scores (LOD score) are aligned below the effects. Effects pattern matches the distribution of rs262996903 alleles among the J:DO founder strains. (C) Haplotype effects for CES1C *cis* pepQTL for a peptide with LKAEEVAFWTELLAK sequence. Association scores (LOD score) are aligned below the effects. Effects pattern matches the distribution of rs31983697 alleles among the J:DO founder strains. (D) Amino acid sequence for CES1C (P23953). Peptide sequences are highlighted, and corresponding coding variants are in bold text. QKTESELLEISGK (blue) has an arginine (R) polymorphic site immediately upstream of it. For mice lacking the R residue (rs262996903), a trypsin cleavage site, no QKTESELLEISGK peptide will be produced from trypsin digest. LKAEEVAFWTELLAK (green) contains the coding variant (rs31983697).

**Figure S2.**
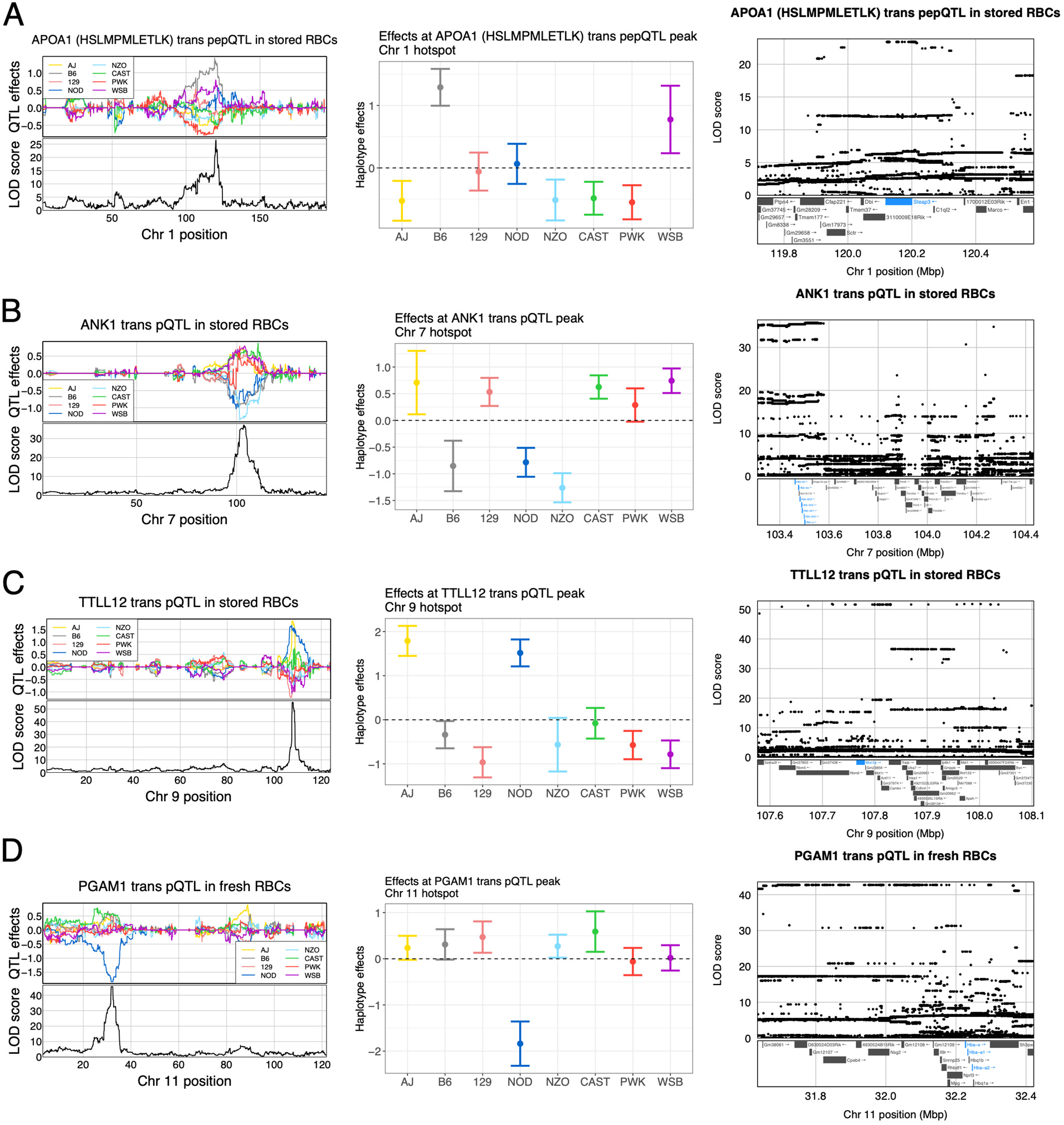
Haplotype effects and SNP associations at for example QTL at the four QTL hotspots. (A) APOA1 maps a *trans* pepQTL to the chromosome 1 hotspot in stored RBCs. (Left) Haplotype effects for APOA1 *trans* pepQTL that maps to the *Steap3* locus. Association scores (LOD score) are aligned below the effects. (Middle) Haplotype effects at the peak locus with standard error bars. (Right) SNP associations within the QTL region aligned with encoded genes. *Steap3* is highlighted. (B) ANK1 maps a *trans* pQTL to the chromosome 7 hotspot in stored RBCs. (Left) Haplotype effects for ANK1 *trans* pQTL that maps to the *Hbb* locus. Association scores (LOD score) are aligned below the effects. (Middle) Haplotype effects at the peak locus with standard error bars. (Right) SNP associations within the QTL region aligned with encoded genes. *Hbb* gene cluster is highlighted. Olfactory genes were removed to improve clarity. (C) TTLL12 maps a *trans* pQTL to the chromosome 9 hotspot in stored RBCs. (Left) Haplotype effects for TTLL12 *trans* pQTL that maps to the *Mon1a* locus. Association scores (LOD score) are aligned below the effects. (Middle) Haplotype effects at the peak locus with standard error bars. (Right) SNP associations within the QTL region aligned with encoded genes. *Mon1a* is highlighted. (D) PGAM1 maps a *trans* pQTL to the chromosome 11 hotspot in fresh RBCs. (Left) Haplotype effects for PGAM1 *trans* pQTL that maps to the *Hba* locus. Association scores (LOD score) are aligned below the effects. (Middle) Haplotype effects at the peak locus with standard error bars. (Right) SNP associations within the QTL region aligned with encoded genes. *Hba* gene cluster is highlighted.

**Figure S3.**
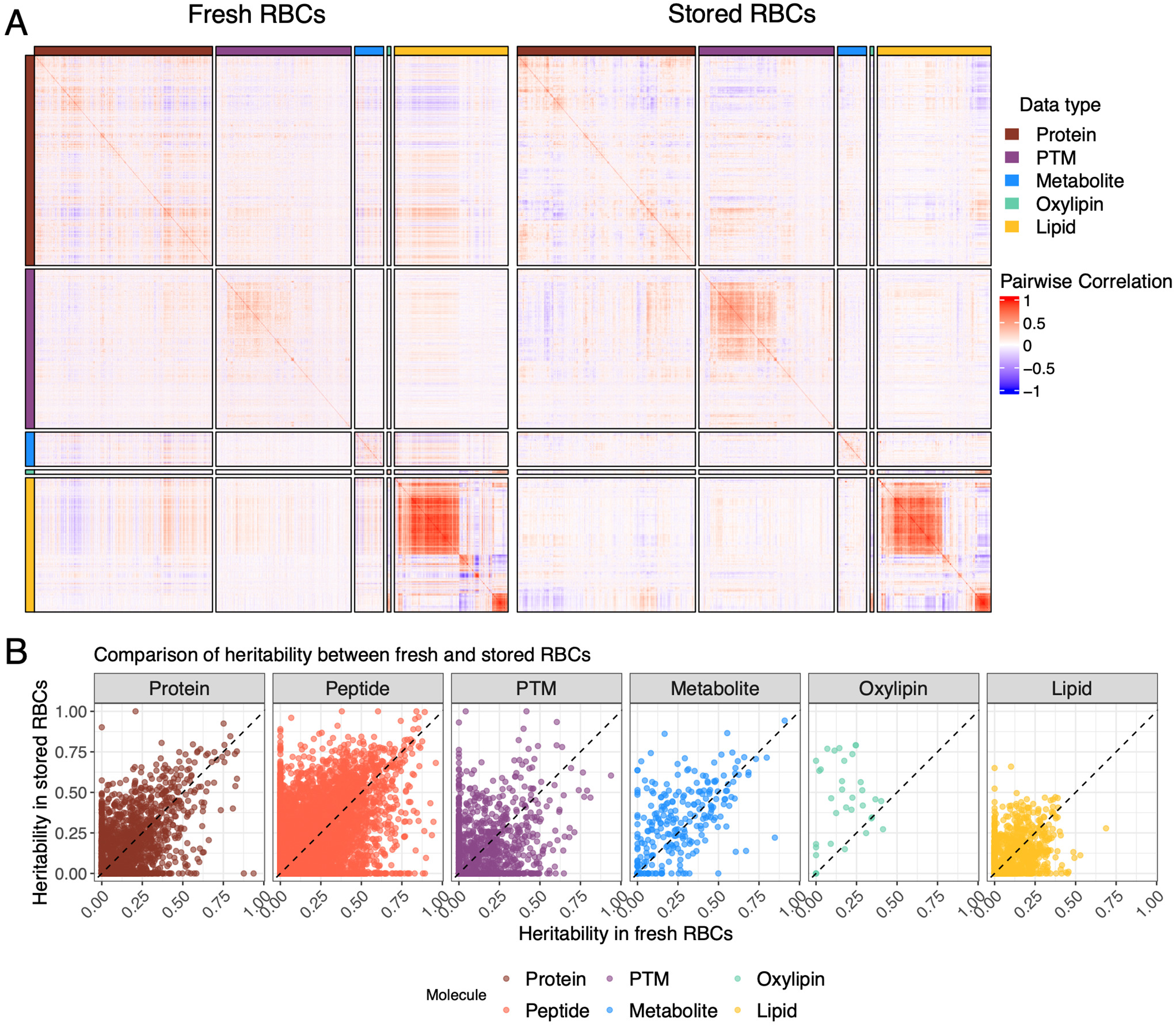
Correlation patterns and heritability across RBC omic data. (A) Heatmap of the pairwise correlations between abundance levels for proteins, PTMs, metabolites, oxylipins, and lipids in fresh (left) and stored (right) RBCs. Peptide data layer excluded for clarity due to overwhelming size. (B) Comparison of heritability for proteins, peptides, PTMs, metabolites, oxylipins, and lipids between fresh and stored RBCs. Diagonal black dashed lines included for reference.

**Figure S4.**
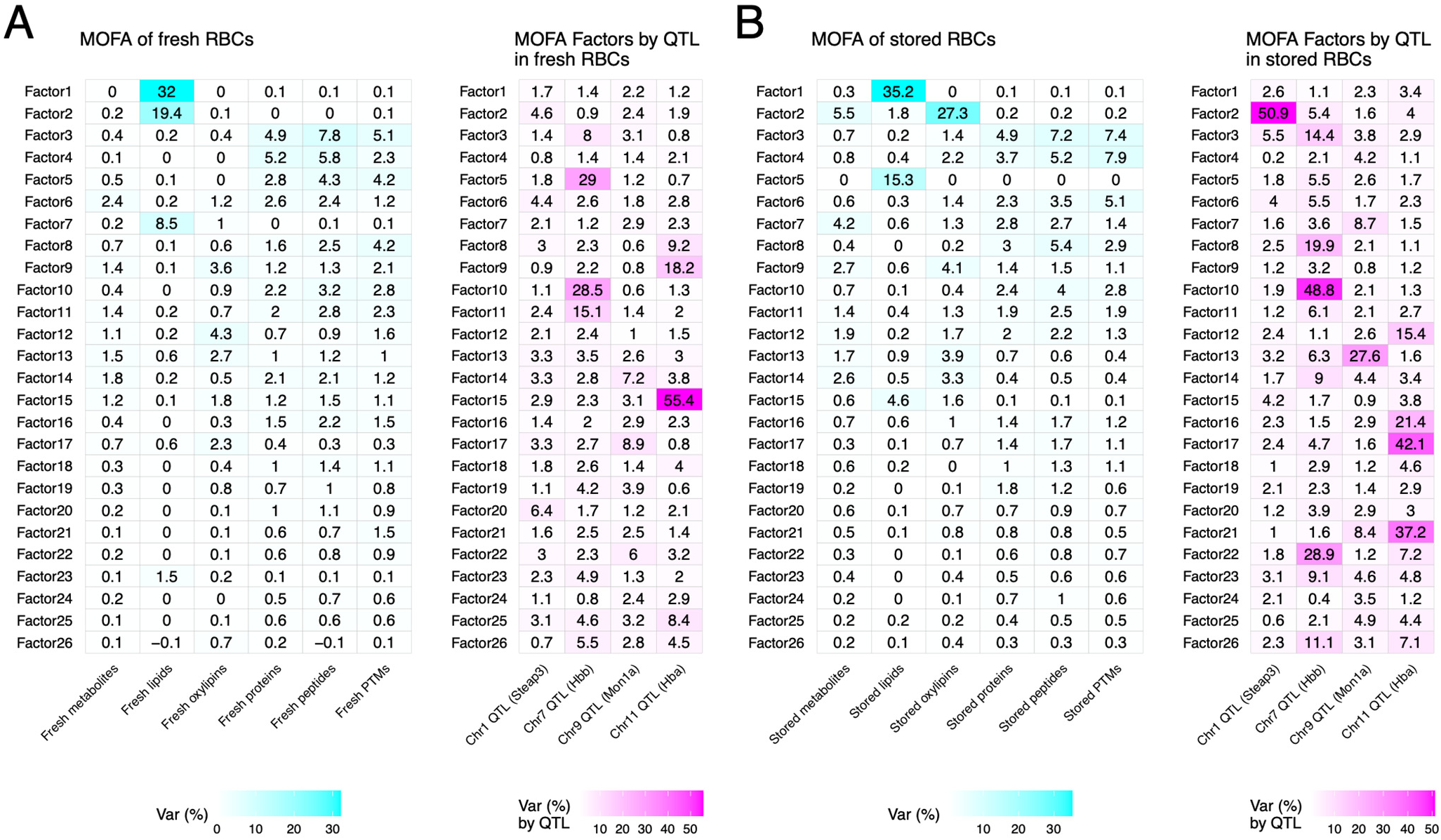
Multi-omics factor analysis (MOFA) of full RBC data. Heatmaps of the proportion of variance from MOFA for the (A) full fresh RBC omic data and the (B) full stored RBC omic data. Data layers comprised proteins, peptides, PTMs, metabolites, oxylipins, and lipids. Variance explained for factors by data layer (left) and for factors by hotspot locus (right).

**Figure S5.**
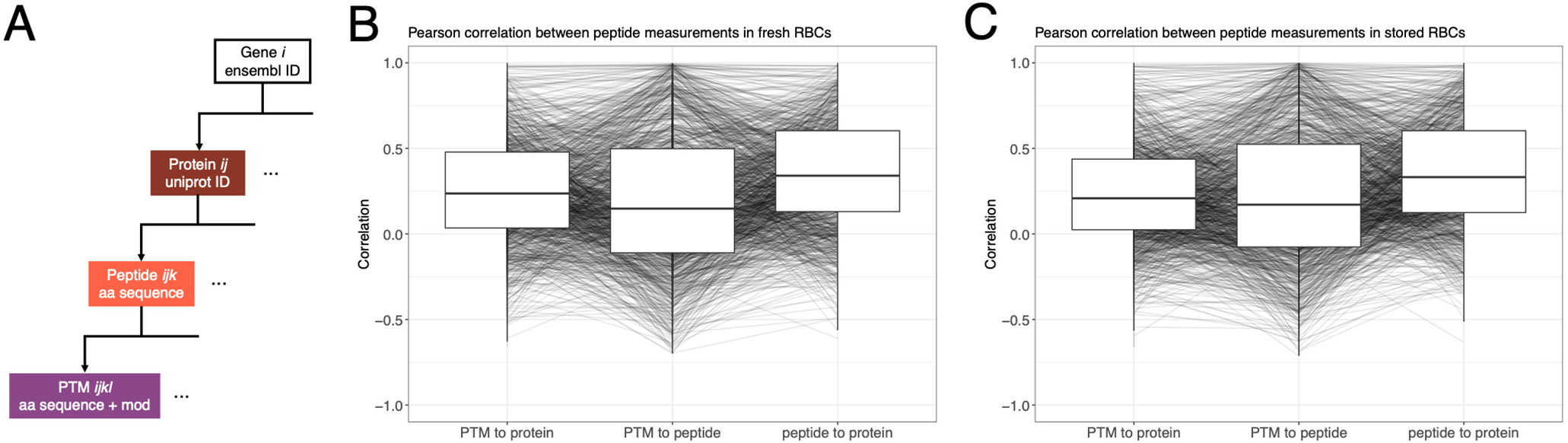
Relationships between protein, peptide, and PTM measurements. (A) Diagram of the hierarchy of the proteomics data. A gene may map to many proteins, which may map to many peptides, which may map to many PTMs. Boxplots of the pairwise correlations between measurements linked by peptide sequence per uniprot ID in (B) fresh and (C) stored RBCs. PTMs were unadjusted for peptides for this analysis.

**Figure S6.**
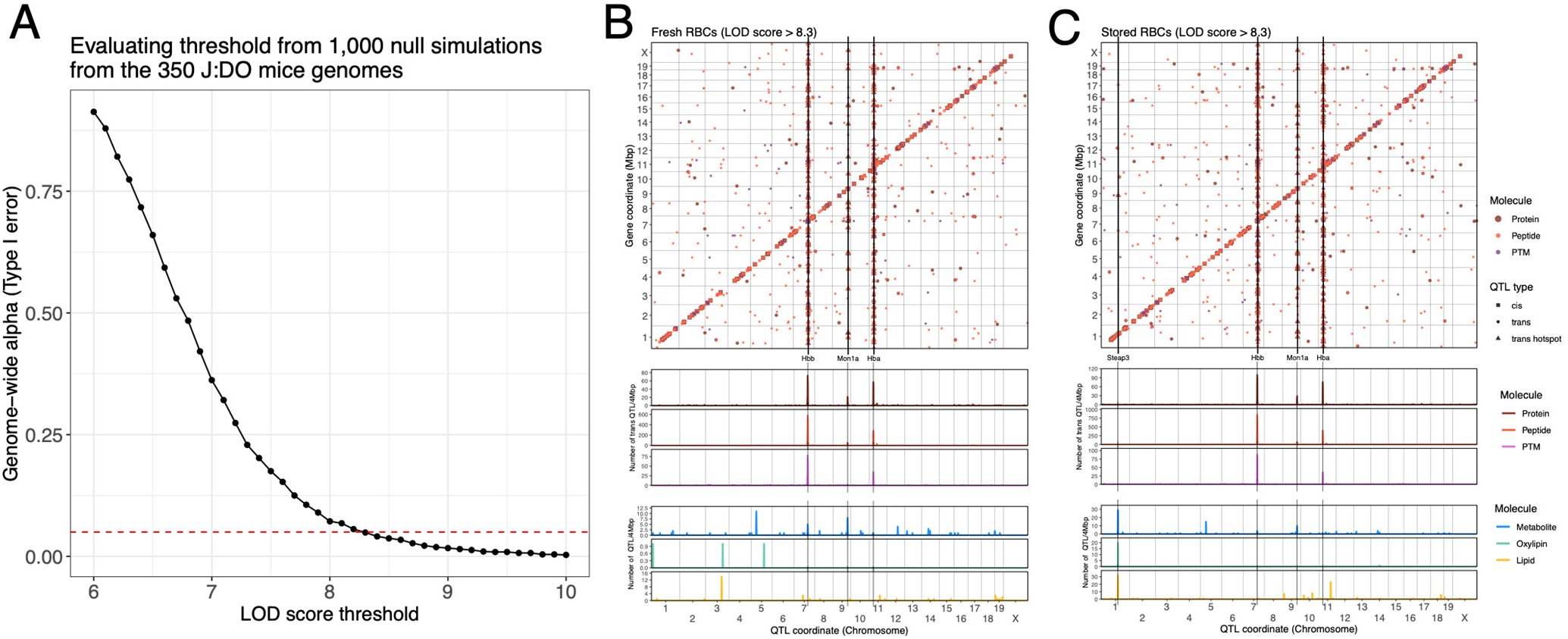
Simulation-based genome-wide type I error rates for QTL mapping. (A) Type I error rate by LOD score threshold from 1,000 simulations based on the 350 J:DO mice genomes used in this study. Horizontal red dashed line represents a genome-wide type I error rate of 0.05. QTL maps (LOD score > 8.3) for proteins, peptides, and PTMs in (B) fresh and (C) stored RBCs, aligned with QTL densities (based on a 4 Mbp window) for proteins, peptides, PTMs, metabolites, oxylipins, and lipids. Shape of data point indicates type of QTL (*e.g.*, *cis, trans,* hotspot). Vertical lines included to highlight QTL hotspots at the *Steap3*, *Hbb*, *Mon1a*, and *Hba* loci. More lenient QTL maps (LOD score > 7) are available in Figure 2A-B.

## SUPPLEMENTARY TABLES

**Table S1.**
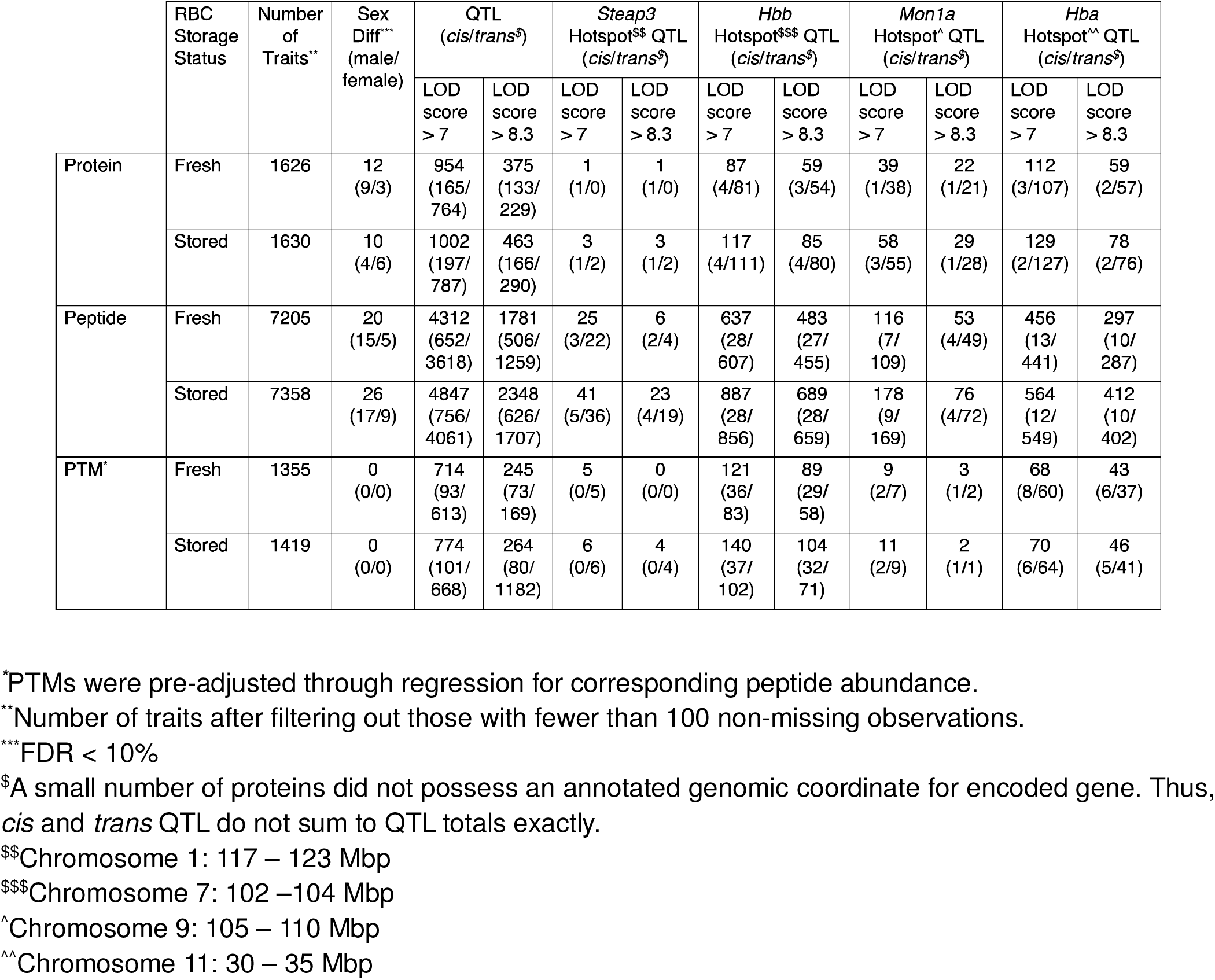
Detected sex differences and QTL for proteins, peptides, and PTMs.

## SUPPLEMENTARY FILES

**Supplementary Data File 1. QTL results for proteins, peptides, and PTMs in fresh and stored RBCs.**

